# Repeats Mimic Pathogen-Associated Patterns Across a Vast Evolutionary Landscape

**DOI:** 10.1101/2021.11.04.467016

**Authors:** Petr Šulc, Andrea Di Gioacchino, Alexander Solovyov, Sajid A. Marhon, Siyu Sun, Håvard T Lindholm, Raymond Chen, Amir Hosseini, Hua Jiang, Bao-Han Ly, Parinaz Mehdipour, Omar Abdel-Wahab, Nicolas Vabret, John LaCava, Daniel D. De Carvalho, Rémi Monasson, Simona Cocco, Benjamin D. Greenbaum

## Abstract

An emerging hallmark across human diseases – such as cancer, autoimmune and neurodegenerative disorders – is the aberrant transcription of typically silenced repetitive elements. Once active, a subset of repeats may be capable of “viral mimicry”: the display of pathogen-associated molecular patterns (PAMPs) that can, in principle, bind pattern recognition receptors (PRRs) of the innate immune system and trigger inflammation. Yet how to quantify the landscape of viral mimicry and how it is shaped by natural selection remains a critical gap in our understanding of both genome evolution and the immunological basis of disease. We propose a theoretical framework to quantify selective forces on virus-like features as the entropic cost a sequence pays to hold a non-self PAMP and show our approach can predict classes of viral-mimicry within the human genome and across eukaryotes. We quantify the breadth and conservation of viral mimicry across multiple species for the first time and integrate selective forces into predictive evolutionary models. We show HSATII and intact LINE-1 (L1) are under selection to maintain CpG motifs, and specific Alu families likewise maintain the proximal presence of inverted copies to form double-stranded RNA (dsRNA). We validate our approach by predicting high CpG L1 ligands of L1 proteins and the innate receptor *ZCCHC3*, and dsRNA present both intracellularly and as MDA5 ligands. We conclude viral mimicry is a general evolutionary mechanism whereby genomes co-opt pathogen-associated features generated by prone repetitive sequences, likely offering an advantage as a quality control system against transcriptional dysregulation.

## MAIN TEXT

It has recently become clear that repetitive elements, which represent most of the human genome and often derive from integrated viruses and genome parasites, can function as “non-self” pathogen-associated molecular patterns (PAMPs). Under aberrant conditions, such as cancer^1^ and viral infection^2–4^, repeats may be overexpressed, where the PAMPs they display can engage the innate immune system^5–10^. Consistently, a growing body of literature has demonstrated aberrant expression of immunostimulatory repeats across an array of human diseases, such as in aging^11^ and autoimmunity^12^, implying viral mimicry may be a fundamental feature across inflammatory diseases. The ability to quantify PAMPs capable of being sensed by pattern recognition receptors (PRRs) is also of considerable theoretical interest^13^. Mathematical models of the evolution of human H1N1 influenza since the 1918 pandemic showed an attenuation of CpG motifs, leading to the prediction such motifs can be targeted by PRRs^14, 15^. It was subsequently discovered the protein ZAP (*ZC3HAV1*) is a PRR targeting CpG motifs, indicating inferences drawn from genome evolution can predict PAMPs and PRR specificities relevant to the adaptation of emerging viruses^16, 17^, including SARS-CoV-2^18, 19^. It has been more difficult to predict specificities associated with structure formation^20^, such as the recognition of double-stranded RNA PAMPs by MDA5 (*IFIH1*) or TLR3^21^, or other more complex PAMPs, such as the creation of DNA:RNA hybrids. Importantly, viral mimicry can be leveraged therapeutically: the expression of immunostimulatory repeats is inducible by epigenetic drugs, leading to triggering of innate immune sensors and induction of an interferon response^5–9^.

Several fundamental questions remain, such as which PRRs can be activated by which specific repeats, if viral mimicry serves a functional role in the genome as an evolved checkpoint for loss of epigenetic regulation or genome fidelity, and whether tumors and pathogens adapt to manipulate mimicry to their own selective advantage^22, 23^. In one evolutionary scenario, repeats which contain PAMPs in somatically silenced regions can offer a fitness advantage to cells due to their ability to trigger PRRs under epigenetic stress, eliminating dysregulated cells and maintaining tissue homeostasis^22, 23^. Such features could then be maintained by natural selection. Alternatively, in a neutral scenario, it may be that high RNA concentration resulting from transcriptional dysregulation can engage PAMPs non-specifically, and their sensing is a by-product of dysregulation rather than of selection acting on specific features. Discriminating between such scenarios is critical to understanding how non-self mimicry by the self-genome evolved, and how it can be leveraged for emerging therapies and honed for existing ones. There is therefore a pressing need for computational approaches to quantify the presence of viral mimics, define their immunological features and quantify the evolutionary dynamics of their (putative) PAMP content. We utilize a novel approach from statistical physics to quantify nucleic-acid motifs and double-stranded structures under selective forces and use multiple assays to validate our predictions. In doing so we define specific categories of repeat families that were likely retained by natural selection to display viral mimics.

### Inference and evolutionary dynamics of pathogen-associated patterns in repeats

We utilize the framework of selective and entropic forces to infer the presence and evolutionary dynamics of a PAMP or set of PAMPs in a genome^14^. Sequences incorporated into a genome, subject to constraints such as local nucleic acid content, accumulate mutations during evolution to resemble, on average, the self-genome while selective forces, such as those acting on an atypical “non-self” pattern, oppose such a trend (**Fig. 1A**). The selective force is an intensive parameter that can be interpreted as a measure of the depletion (negative) or excess (positive) of a feature in a genome sequence beyond the degree it would be expected based on the nucleotide statistics within the sequence. This framework is ideal for the study of viral mimicry and PAMP detection as selective forces can be readily compared between groups of sequences independently of their length. We infer selective forces on a given sequence for one or more patterns introducing them as parameters in a Maximum Entropy model for genomic sequences^24^ (Methods). Our inference algorithm uses exact transfer matrix methods from statistical physics which, unlike earlier approaches^15^, are computationally efficient (scaling with the length of the sequence) and facilitate the analysis of longer sequences and large sequence datasets (Methods).

**Figure 1.**
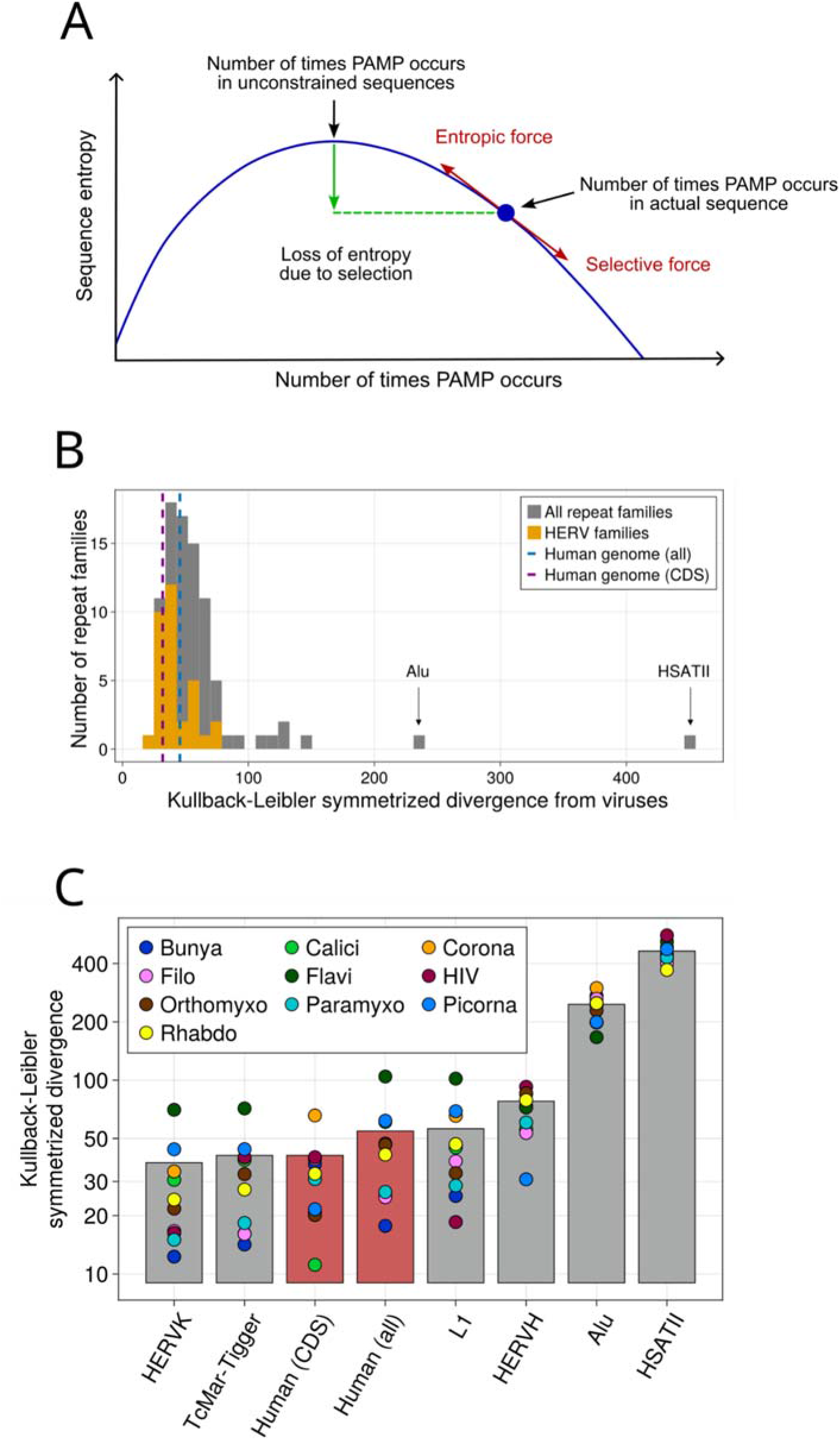
Competition between selective and entropic forces define presence of pathogen associated patterns in the genome. **A,** Representation of selective versus entropic forces on a PAMP. Random sequences reproducing the nucleotide frequencies generically have numbers of occurrences of a PAMP significantly different from what is observed in an actual genomic sequence. Deviations imply constraints to enrich or avoid a PAMP, which we characterized by a, respectively, positive or negative selective force. In our Maximum Entropy framework, the selective force counterbalances the entropic force resulting from the loss of diversity (entropy) in sequences having statistically abnormal PAMP numbers. **B,** Comparison of all nucleotide biases between repeat and viral families. In this histogram each repeat family is compared with reference viral genomes of viral families by computing the symmetrized Kullback-Leibler divergence of the probability distributions associated to models inferred from viral and repeat sequences. Repeats with particularly high divergence values are indicated by arrows. The value “Human genome (all)” is obtained with a model trained on sequences randomly sampled from the human genome (hg38), while for “Human genome (CDS)” we only consider coding sequences. **C,** Detailed comparison of nucleotide biases between selected repeat and viral families. Each point is the symmetrized Kullback-Leibler divergence between a repeat family (x-axis) and a specific viral family (as indicated by the color). Bars represent, for each repeat family under consideration, the average value over the viral families.

A repetitive element is primarily defined by the presence of multiple copies (inserts) of its sequence in the genome. As additional repetitive copies accumulate, we measure the evolutionary dynamics of PAMPs as they diverge from their original sequence. We use two approaches. The first uses relaxation dynamics: a new repeat in a genome evolves until it reaches a genomic equilibrium value determined by a balance of factors such as constraints on nucleic acid usage and forces on sequence patterns. By analogy, selective forces drove the avian origin 1918 influenza virus towards a new equilibrium in humans with lower CpG content, and it was subsequently found the PRR ZAP consistently targets CpG with greater affinity in humans than birds^25^. The second approach uses a Kimura-based model as a proxy for the neutral evolution of a sequence with given PAMP content. We implement this variant of the Kimura model numerically to provide a null model of repeat evolution within a genome. As in the standard Kimura model, we use different mutation probabilities for transitions (a purine mutating into a purine or a pyrimidine into a pyrimidine) and transversions (a purine mutating into a pyrimidine or vice-versa), with the former being more probable than the latter (see Methods). Additionally, we use different ratios of mutation rates corresponding to nucleotide transitions and transversions in CpG and non-CpG context^26^. We calculated the dinucleotide distribution stationary value, obtained as the stationary vector of the stochastic matrix with entries corresponding to probabilities of mutating from one dinucleotide to another dinucleotide (see Methods and Table 1 therein).

To test our approach, we quantify the overall degree of motif usage similarity between families of human infecting viruses and regions of the human genome. We infer Maximum Entropy models with forces on all single, di-, and tri-nucleotide motifs for a set of human repeat families and compare them to models inferred for families of viruses which infect humans (**Fig. 1B, C**). To quantify similarity of motif usage in the two sets of families we use the symmetrized Kullback-Leibler divergence (details about its computation are given in Methods) between the corresponding models. Primarily viral and human genomes share similar overall motif usage, a form of mimicry that is likely a product of shared constraints on nucleotide usage across organisms and viruses, with some minor variation between viral families. Coding regions in the human genome show stronger overall similarity to human infecting viruses, most of whose genomes are devoted to coding, than non-coding regions, although large variation exists in the latter. For instance, consistent with the overall trend, HERVK repeats show the strongest similarity with viruses among repeat regions. As a stark exception, we find far less motif usage similarity between Alu repeats and HSATII than either to the rest of the human genome or to human infecting viruses. Neither repeats encode known proteins and both are thought to have non-viral origins^27^, indicating such regions may be subject to different evolutionary pressures from the other repeats considered here.

### Landscape of repeats with selective forces on CpG dinucleotides

CpG dinucleotides in humans are PAMPs in DNA, recognized via TLR9^28^, and, as has been seen more recently, CpGs in RNA are engaged via ZAP^17^. We compare the evolution of individual dinucleotide motifs (quantified by calculating the selective force, *x_m_*, on a dinucleotide motif, *m*, as defined in Methods) between the original consensus sequence, representing the sequence most likely to resemble a founding ancestral repeat insertion, and its subsequent copies in the genome (**Fig. 2**). We analyzed *x*_CpG_, and all other *x_m_*, for all repeat families annotated in the DFAM database^29^, finding outliers such as Alu repeats and HSATII, the latter consistent with previous results^10^. Typically, CpG content in the human genome is highly underrepresented as CpG sites mutate at a much faster rate than the rest of the genome^26, 30, 31^. We plot the mean difference in *x*_CpG_ per repeat family versus *x*_CpG_ for the consensus insert (**Fig. 2A**). Consistent with our assumptions, we see families where the selective force on CpG dinucleotides for the progenitor insert was greater than −1.9 have decreased their force to this value, while those less than −1.9 have increased their value. We therefore establish a genome-wide equilibrium in line with equilibria observed for human adapted viruses such as influenza^15, 18^. If a repeat is not subject to selection, one would expect its insertions to evolve according to a Kimura model with respective mutation rates for transitions and transversions, an approach used in sequence evolution models to explain lower CpG content in vertebrate genomes^32–34^. **Fig. 2B** shows the relaxation of *x*_CpG_ as a function of the Kimura distance^35^ used for each individual repeat sequence, as a proxy for time since insertion. The Kimura distance is the expected number of mutations accumulated in a given period of time by a sequence that evolves with a higher probability of transitions over transversions. It represents the expected number of differences between two sequences after a given period of time at fixed mutational rates. We use it as a proxy for time since insertion for each individual repeat sequence, relative to other elements of the same family. Most repeat families show relaxation to the mean genome force expected from the neutral model, further implying HSATII may be specifically under selection to hold this PAMP. Moreover, HSATII is the most represented repeat among those overlapping with high-*x*_CpG_ (*x*_CpG_ > 0) genomic regions in the human genome (**Fig 2C**).

**Figure 2.**
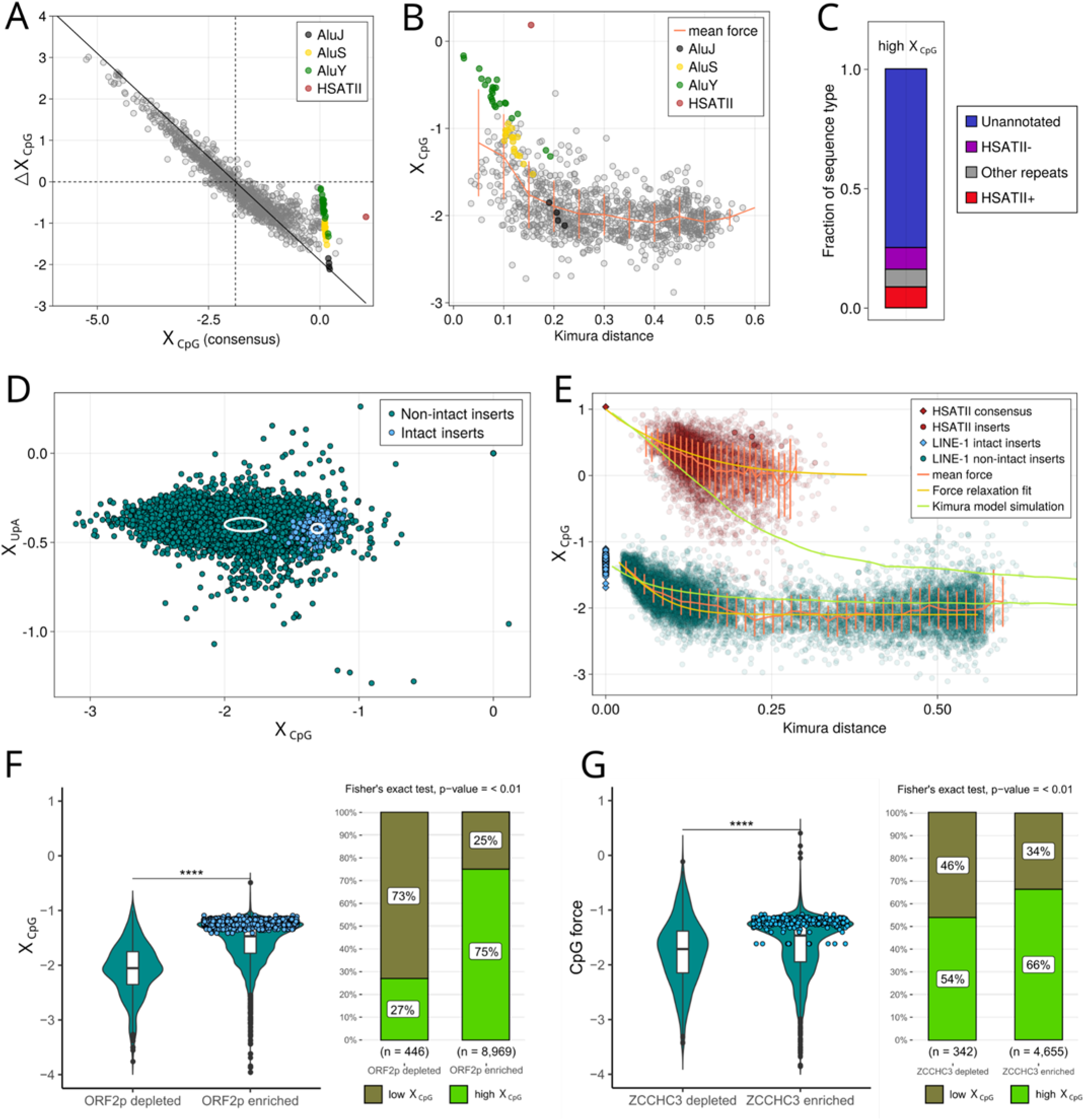
Forces on CpG dinucleotides in the human genome. **A,** Change in *x*_CpG_ computed on all inserts annotated in hg38 for each repeat family, versus *x*_CpG_ of the consensus repeat reported in the DFAM database. Alus and HSATII are highlighted as exceptions to the general trend. **B,** The mean *x*_CpG_ of all inserts in a repeat family as a function of the Kimura distance from the consensus sequence for each family. **C,** Annotation of high-*x*_CpG_ (*x*_CpG_ > 0) sequences in the human genome according to their overlap with annotated repeats in the DFAM database. The + or - sign after the repeat name indicates the sense in which the repeat is annotated in the database. “Unannotated” sequences do not overlap with any repeat in the database. **D,** Scatter plot of *x*_CpG_ and *x*_UpA_ for LINE- 1 functional (blue) and non-functional (green) elements in the human genome. The white ellipse corresponds to one standard deviation distance from the mean for *x*_CpG_ and *x*_UpA_ forces on FLI and FLnI LINE-1 inserts respectively. **E,** *x*_CpG_ for FLnI inserts of LINE-1 and HSAT-II in human genome as a function of average distance from the intact FLI sequences (for LINE-1) or the distance from the consensus sequence (for HSAT-II). The force relaxation evolutionary model fit is shown for both sequence families together with a Kimura (null) model fit. **F,** Distribution of *x*_CpG_ of L1 ORF2p binding L1 transcripts in embryonal carcinoma cell line (N2102Ep). Functional intact LINEs are colored in blue (BH corrected p-value labeled for t-test, **** denotes adjusted p-value < 0.01). ORF2p enriched and depleted transcripts are selected by differential expression analysis between ORF2p-IP versus Mock/total with |log2FC| greater than 3 and adjusted p-value < 0.05 for Fisher Exact test on proportion of *x*_CpG_ high versus *x*_CpG_ low of ORF2p enriched and depleted transcripts. **G,** *x*_CpG_ on *ZCCHC3* binding LINE transcripts in N2102Ep. Functional intact LINEs colored in blue (BH corrected p-value labeled for t-test, **** denotes adjusted p-value < 0.01). *ZCCHC3* enriched and depleted transcripts selected by differential expression analysis between *ZCCHC3*-IP versus Mock/total with a |log2FC| greater than 3 and adjusted p-value < 0.05.

As L1 elements have the most copies in the genome, they are most amenable to our approach. Their copies are estimated to constitute about 20% of human genome^36^. Here we only consider full-length inserts, as annotated in L1Base2, and contrast those designated as fully intact (denoted FLI), from full-length sequences designated as non-intact (FLnI)^37^. Fully functional L1 DNA sequences are regulated by hyper-methylation at CpG sites in their promoter, to inhibit their transcription^38, 39^. Indeed, we find FLI L1 have higher CpG content than FLnI (**Fig. 2D**), though most conserved CpGs are not in the promoter region (**Supplementary Fig. S1**). We find that as a L1 genome insertion ceases to contain an intact copy, its CpG content decays with the Kimura distance to the consensus, reaching the genome mean in a predictable way according to the Kimura model for neutral genomic evolution (**Fig. 2E**). The most recent inserts into the human genome therefore appear to not have equilibrated. It is important to identify all such cases because families that have not saturated are candidates for viral mimicry via PAMP display, such as when LINE-1 is overexpressed in tumors^1, 40–42^. For Alu repeats we observe a pattern of CpG-content relaxation similar to LINE-1, but only when considering together the major Alu subfamilies (AluY, AluS, AluJ). The younger AluY and, to a lesser extent, AluS are not yet equilibrated and still possess PAMP-like high CpG content. (**Extended Data Fig. 1A)**. For HSATII, evolutionary dynamics of the force relaxation (**Fig. 2E**) corresponds to saturation at force approximately equal to −0.4, well above the equilibrium distribution computed from the Kimura model, implying its ability to retain CpGs is maintained by selection. For most families the data points are scarce and noisy, making a relaxation fit such as the one shown for HSATII and LINE-1 difficult. **Supplementary Table 1** lists the full repeat atlas of CpG content, computed both for the consensus repeat and as an average over the inserts in the genome. CpG-rich regions (*x*_CpG_ > 0) in the human genome mostly concentrate in intergenic and, to a lesser extent, intronic regions (**Extended Data Fig. 1B**), and are listed in **Supplementary Table 2**.

We reasoned that the force acting on CpGs in intact L1 species is enforced by the *in cis* binding of L1 encoded proteins. To determine if this is the case, we analyzed the RNAs affiliated with both L1 ORF1p and ORF2p by RNA co-immunoprecipitation sequencing (RIP-seq). Notably ORF2p is the reverse transcriptase of L1. We conducted α-ORF1p RIP-seq in N2102Ep human embryonal carcinoma cell lines^43^ and, for the first time, α-ORF2p RIP-seq. Transcripts enriched by co-IP were determined by differential expression analysis of RIP versus total RNA and matched mock IP controls (Methods). Intact L1s are exclusively recovered in ORF1p and ORF2p binding transcripts (**Fig. 2F; Extended Data Fig. 1C**) as expected. Consistently, ORF2p (**Fig. 2F**) and ORF1p (**Extended Data Fig. 1C**) enriched transcripts (Log2FC > 3, adj. p-val. < 0.05) have CpG forces consistent with the high *x*_CpG_ observed in fully intact transcripts and compared to controls (Log2FC < -3, adj. p-val. < 0.05). To further examine if innate immune receptors co-IP with high *x*_CpG_ L1 RNA, we examined the ligands of *ZCCHC3*, a protein recently described as a co-sensor of cGAS^44^ that has been found to interact with L1 ribonucleoprotein in an RNA-dependent fashion^45, 46^. We find a substantial enrichment in high CpG L1 RNA associating *ZCCHC3*, compared to both controls and to non-intact L1 (**Fig. 2G**). Our findings indicate high *x*_CpG_ L1 RNA is both more likely to associate with L1 proteins and with a putative innate immune sensor of L1 RNA. We therefore conclude high *x*_CpG_ is associated with both replication competent L1 and innate immune sensing of L1.

### Landscape and evolution of repeats with selective forces on double-stranded RNA formation

We further extend our approach, for the first time, to the formation of anomalous secondary structure by calculating the force on double-stranded RNA (dsRNA) formation in repeats (**Fig. 3**, **Extended Data Fig. 2**). This value quantifies the tendency of an RNA transcript to form double-stranded segments. It is generally accepted that Toll-Like Receptor 3 (TLR3) is activated by short (approx. 30 bp) endosomal dsRNA and Retinoic acid-Inducible Gene I (RIG-I, *DDX58*) by short (tens of bases) cytoplasmic dsRNA accompanied by a triphoshphate^40^, while Melanoma Differentiation Associated protein 5 (MDA5) recognizes longer cytoplasmic dsRNA associated with RNA virus replication^47^. We calculated the double stranded force, *x*_ds_ (Methods), for repetitive families as well as ncRNA and mRNA sequences in the human genome, and randomly generated sequences (**Fig. 3A**). While the mean value of *x*_ds_ computed for functional mRNA sequences and noncoding sequences is close to zero and essentially the same as the value for random sequences, the consensus sequences of repeats contain multiple families with long complementary segments contributing to an increased average *x*_ds_ value (34 families out of 980 analyzed have *x*_ds_ > 0,5). While the general trend is to relax *x*_ds_ towards zero (**Fig. 3B**), we observe outliers having a higher positive *x*_ds_ value, indicating a possible reservoir of double-stranded segments being maintained by selection. Including are the DNA transposons Tigger4a (**Extended Data Fig. 2B**), MER107 and MER6B (**Fig. 3B**), which could be transcribed under aberrant conditions.

**Figure 3.**
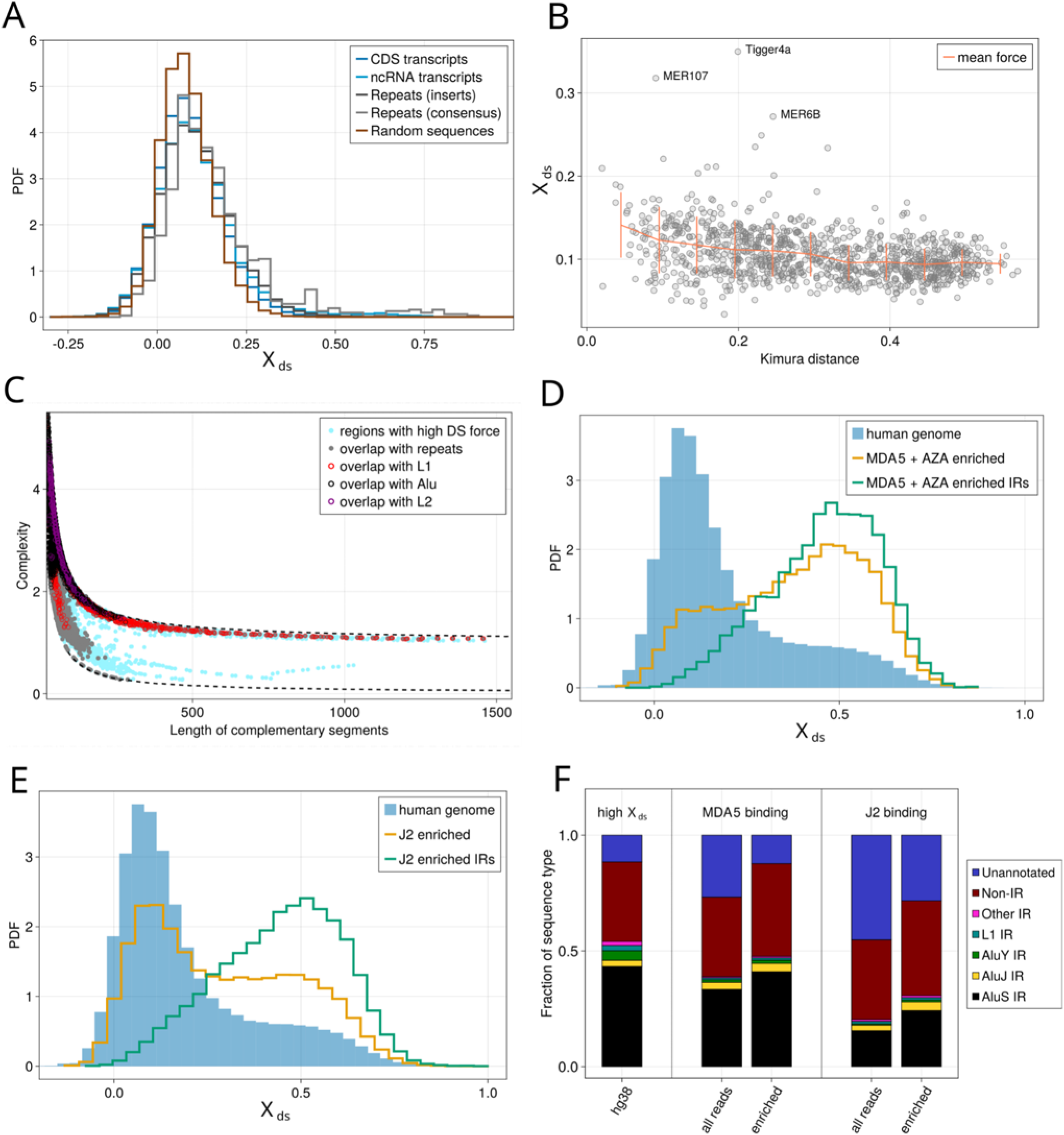
Double-stranded forces in the human genome. **A,** Histogram of *x*_ds_ calculated for mRNA coding sequences, non-coding RNAs, inserts, consensus sequences of repeats, and sequences obtained by randomly reshuffling mRNA coding sequences (yellow). **B,** Mean of *x*_ds_ calculated for each family of repeats as a function of the mean Kimura distance of all inserts in a repeat family from their consensus sequence. The solid line corresponds to mean value (and standard deviation from it) for all families binned into the same distance from consensus. **C,** Complexity of sequences in complementary regions found in the human genome as a function of segment length. Complementary regions that overlap with known repeat element or ncRNA or mRNA are highlighted as gray dots, with different contour colors depending on the specific family they overlap with. Dashed lines correspond to the complexity of a completely random sequence (top line) and trivial region consisting of a single nucleotide (bottom). Complexity of both complementary segments are similar, so we only include the complexity of one of each complementary transcript. **D,** *x*_ds_ histograms in human genome (sliding window with transcript of length of 3 kb) compared to MDA5 binding RNA transcripts. Enriched transcripts have a positive log-enrichment with respect to the control experiment. Inverted repeat (IR) transcripts are annotated repeats with another repeat of the same family in opposite genomic sense within 3 kb. **E,** Similar to panel **(D)**, for J2 binding transcripts. **F,** Type of repeat (as annotated in RepeatMasker) with the longest overlapping sequence in complementary sequences for high-*x*_ds_ (*x*_ds_ > 0.5) windows in hg38 (left), the MDA5 binding experiments (middle) and the J2 binding experiment (right). Sequences are accounted as “IR” (Inverted Repeats) if the two complementary regions overlap with repeats annotated in the database with the same name but inverted sense (+/- or -/+). “Non-IR” indicates cases where the two repeats overlapping with the two complementary regions have a different name. “Unannotated” indicates cases where one or both the two complementary regions do not overlap with any repeat in the database.

To locate possible sources of double-stranded segments originating from the same transcript, we scan the entire genome (hg38 assembly), using a window of transcripts of length 3000bp, comparable to typical lengths of long ncRNAs^48^. We quantified the sequence complexity of such complementary segments (based on Kolmogorov complexity, Methods), as shown in **Fig. 3C**. The segments close to the low complexity limit typically contain a repeating motif of only a few nucleic acids (such as poly(AT)) while the longest segments have higher complexity, i.e. the regions that can form long dsRNA are not exclusively simple repeats (as summarized in an atlas of all families analyzed, **Supplementary Table 2, 3**). We then characterized the distribution of forces in the genomic scan: we observe two peaks, a major one close to 0 and a smaller around 0.5 (**Fig. 3D**). The mean length of the longest complementary segments found in the dataset with *x*_ds_ > 0.5 is 40 base pairs. We found that for the majority (88%) of such regions the complementary segments in the 3000 bases long regions overlap with known repeats. Greater than 43% of identified complementary segments correspond to AluS, where a copy has inserted in a positive orientation close to one in a negative orientation (inverted-repeat Alus, IR-Alus). AluS is the most represented Alu family in the human genome (accounting for more than 60% of the Alu inserts), and it also has the highest fraction of IR inserts, 59% (39% for AluJ and 18% for AluY). In particular, we noticed 73% of the IR-Alus in the human genome consist of AluS IRs, but if we filter for high-*x*_ds_ the share of AluS repeats forming IR- Alus increases to 86% (**Fig. 3F**). Besides Alu subfamilies (which constitute about 50% of long complementary segments that overlap with known inserts), we also identified complementary fragments from the ORF2 open reading frame of LINE-1^49^. We found previously unannotated non-inverted repeats prone to forming long-double stranded RNA (full list in **Supplementary Table 3**). We conclude that while IR-Alus form the major class of binders, other unannotated inverted repeats are also prone to dsRNA formation. Likewise, previous work hypothesized that dsRNA formed from introns is a checkpoint against intron retention^50, 51^. We observed most regions (55.4%) with *x*_ds_ > 0.5 were over-represented at intronic regions (**Extended Data Fig. 2G**).

We next validated our ability to predict double-stranded forming regions. We first examined two published datasets of RNA forming long dsRNA MDA5 receptor ligands, as their transcription has been implicated as a response to genome-wide DNA demethylation^7, 52^. We find that RNA transcripts binding MDA5 under DNA demethylation agents (AZA) display a double-peaked force distribution with a predominance of the large *x*_ds_ peak. Inverted repeats, and notably the AluS family (**Fig 3. D,F**), only populate the high *x*_ds_ peak. AluS repeats account for 89% of enriched IR-Alus in the MDA5-binding experiment, in agreement with our prediction based only on quantifying high-*x*_ds_ sequences in the human genome. A consistent result was found in a second MDA5 ligand dataset (**Extended Data Fig. 2C**^52^). To further validate our ability to predict dsRNA forming transcripts, we generated a novel dataset of sequenced ligands of the J2 monoclonal antibody, an antibody able to recognize dsRNA of greater than 40 bp, nearly identical in length to the average length of anomalous regions predicted when *x*_ds_ > 0.5, in a set of patient-derived colorectal cancer cell lines (Methods, **Extended Data Fig. 2E**). Consistent with our predictions, we show an enrichment of high *x*_ds_ regions in J2 antibody binding transcripts, and with a similar profile as the previously published MDA5 ligands. In this case we found AluS repeats constitute 84% of the enriched IR-Alus, once more in agreement with the value predicted for high-*x*_ds_ sequences with our framework (**Fig. 3E**). These results, based solely on *in silico* analysis of the human genome using our framework, are a striking quantification of the experimental observation that IR-Alus, and especially AluS IR, are the major source of self-RNA that form MDA5 agonists^7^, providing strong validation of the predictive power of our evolutionary model and, in turn, the hypothesis that evolution selected this feature as an epigenetic checkpoint^22, 23^. We further analyzed a dataset of inhibitors of RNA splicing which induce intron retention^53^. We examined RNA sequencing data from SF3B inhibitors which cause the retention of introns in SF3B1 K700E mutant cells. Consistent with our model, we found splicing agents which lead to intron retention over express the high double-stranded force intronic repeats we predicted (**Extended Data Fig. 3, Supplementary Table 5**), supporting the potential ability to manipulate this feature using a cancer therapeutic targeting RNA splicing. Consistently, for inhibitors less associated with intron retention the effect was either weakened or not present. We therefore show a clear ability to predict inverted repeat regions associated dsRNA formation.

### Presence and evolution of PAMPs in genomes across evolutionary scales

To further understand whether PAMPs are held by selection, we examine the presence of repeats with high forces on CpG dinucleotides and double-stranded RNA across 20 genomes (**Extended Data Figs. 4-6**). We calculate the presence of outliers for high *x*_CpG_ and high *x*_ds_ regions across all species. For humans and mouse we show that the presence of such anomalous regions is not primarily due to CpG islands or enhancer regions, based on the FANTOM database^54, 55^ (Methods, **Extended Data Fig. 4**). We find high *x*_ds_ regions occur across many species, even those which do not have the Alu family of SINE elements, providing further evidence such regions are likely a byproduct of the reverse transcription machinery across genomes rather than a function of Alus specifically. To establish such regions in other organisms are not due to low complexity regions, we plot the complexity of high *x*_ds_ regions for the zebrafish genome (**Fig. 4A**). We find many genomic regions which are not low complexity and would be prone to dsRNA formation if transcribed, implying such regions may be a source of PAMPs across species. To the best of our knowledge, this is the first quantification of the presence of likely PAMP-forming repeat regions outside of primates. The full list of regions with *x*_ds_ > 0.5 we discovered in the zebrafish genome is reported in **Supplementary Table 4**.

**Figure 4.**
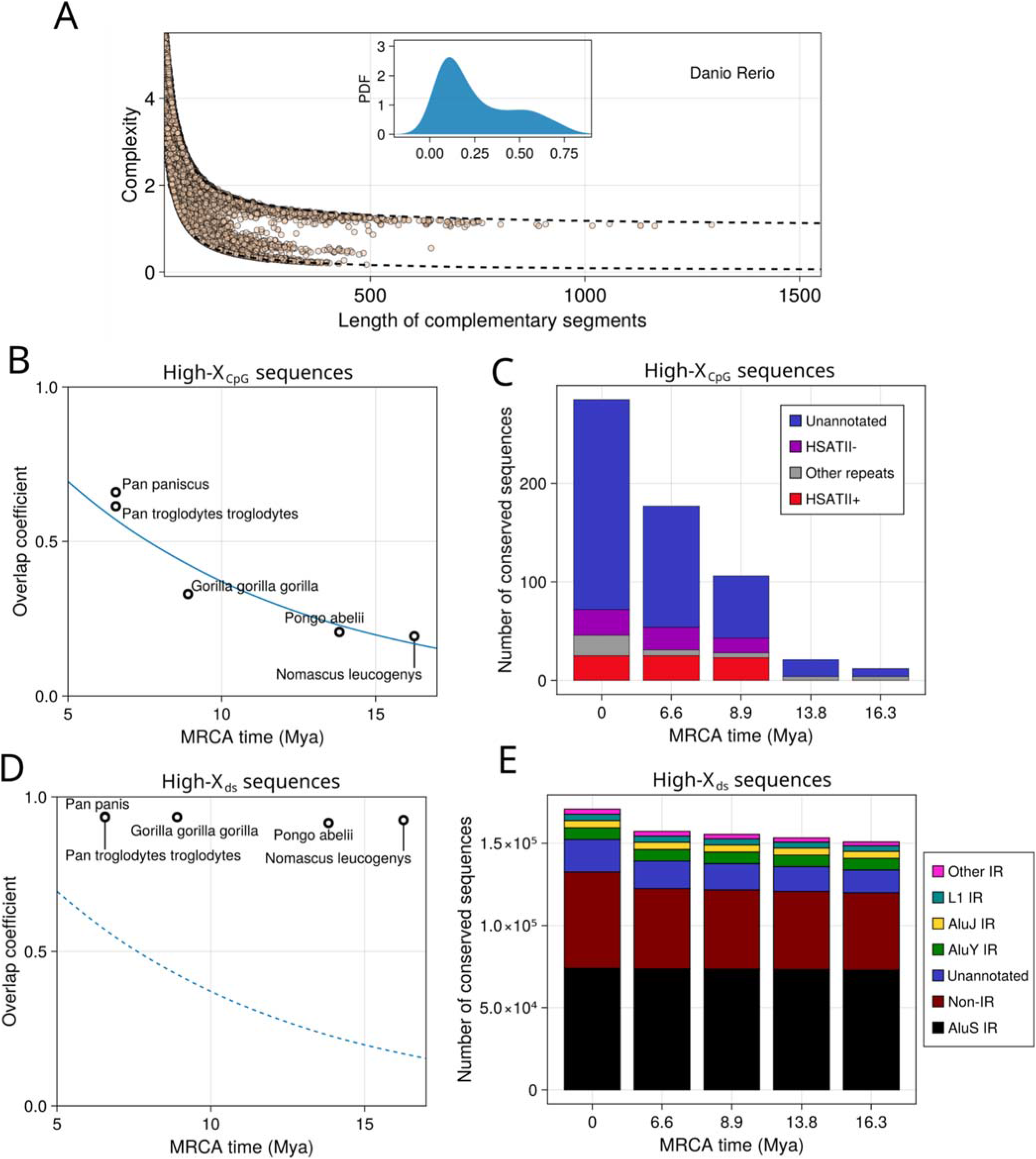
Evolution and conservation of forces on PAMPs. **A,** Complexity of sequences in complementary regions found in the *Danio rerio* genome as a function of segment length. Dashed lines correspond to the complexity of a completely random sequence (top line) and trivial region consisting of a single nucleotide (bottom). **B,** Scatter plot of the overlap coefficient between the high-*x*_CpG_ (*x*_CpG_ > 0) sequences in the human genome and those of other primates versus the most recent common ancestor (MRCA) time^56^. Two high-*x*_CpG_ sequences are considered overlapping if they result as a hit from BLAST (Methods). The blue curve denotes an exponential fit. **C,** Barplot presenting overlap with repeats of conserved high-*x*_CpG_ sequences. The x-axis indicates the MRCA time (0 Mya are human sequences). Sequences are accounted as repeats if they overlap with annotations in the DFAM database. The + or - sign after the repeat name indicates the sense in which the repeat is annotated in the database. “Unannotated” sequences do not overlap with any repeat in the database. **D,** Same analysis as **(C)**, but with high-*x*_ds_ sequences (*x*_ds_ > 0.5). **E,** Barplot presenting overlap with repeats of conserved high-*x*_ds_ sequences. The x-axis indicates the MRCA time (0 Mya are human sequences). Sequences are accounted as repeats if they overlap with annotations in the RepeatMasker database. Sequences are accounted as “IR” (Inverted Repeats) if the two complementary regions overlap with two annotations in the RepeatMasker database with the same name but inverted sense (+/- or -/+). Sequences are indicated as “Non-IR” if the two repeats overlapping with the two complementary regions have a different name. “Unannotated” indicates cases where one or both the two complementary regions do not overlap with any known repeat.

For repeat families identified in humans, we compared selective forces across organisms phylogenetically close to humans. We used the Hominoidea superfamily, whose most recent common ancestor has been proposed to date back to about 16 million years ago^56^. We performed the same analysis as for the human genome, scanning the genomes of five small and great apes and comparing sequences with high *x*_CpG_ and *x*_ds_ values. We first considered the high *x*_CpG_ windows (*x*_CpG_ > 0) and computed the conservation of these sequences across organisms (as quantified by the Overlap Index, Methods). We observed that the number of high *x*_CpG_ sequences conserved between humans and other apes decreases exponentially with their evolutionary distance (**Fig. 4B)**; an expected result, given the high CpG mutation rate. We further observed that, although most high *x*_CpG_ genomic windows do not overlap with any repeat, the vast majority of conserved *x*_CpG_ sequences that do overlap with a repeat are associated with HSATII. HSATII can be found in primates after the branching between the *Pongo* genus and the other great apes, allowing us to pinpoint the HSATII insertion in the primate genomes between 13.8 and 8.9 million years ago. Remarkably, since its insertion into the genome HSATII sequences are conserved to a much greater extent than other sequences in the high-CpG pool (**Fig. 4C**), suggesting a selective pressure maintains PAMPs in HSATII. When we next considered high-*x*_ds_ genomic windows, we found them to be much more generally conserved than high *x*_CpG_ regions (**Fig. 4D**). We found these results striking, since it is not expected by a null model of sequence evolution and implies a selective pressure to keep these windows functionally intact. When focusing on sequences overlapping with repeats, we confirmed inverted Alu repeats are highly conserved in time since their appearance in primate genomes more than 16.3 million year ago (**Fig. 4E**). We therefore conclude Alus, and particularly the AluS family, are likely to have selectively maintained the ability to form double-stranded RNA.

## DISCUSSION

We quantify the landscape and evolutionary dynamics of viral mimicry both across and within genomes. We find, generally, that virus infecting humans mimic the motif usage statistics of human coding regions, indicating shared global constraints on motifs for both viruses and their hosts, consistent with our previous work^14, 15^. There are strong exceptions, such as Alu repeats and HSATII, under less constraint. We find the high-copy satellite RNA HSATII is likely under selection to maintain its pathogen-associated CpG dinucleotides across primates since its origin nearly 10 million years ago and potentially functional L1 inserts maintain atypically high CpG content compared to non-functional copies. We validate the latter with novel co-IPs of high *x*_CpG_ L1 RNA with both L1 ORF1 and ORF2 proteins, indicating such L1s are more likely to be functional, and with the innate immune sensor *ZCCHC3*^44^. Furthermore, we incorporate structure prediction into our method for the first time, which we validate in both published datasets and new dsRNA detecting antibody assays. In humans, many, but not all, dsRNA forming repeats come from inverted Alus, indicating double-stranded RNA, mimicry is likely due to the error prone reverse transcriptional process, rather than being a specific property of the Alus other than their known parasitism of L1. We show a high degree of conservation of double-stranded RNA-forming Alus across primates, indicating selection has maintained their ability to display PAMPs. We find nontrivial potential PAMP forming regions across many genomes which lack either Alus or HSATII, implying reservoirs of potential PAMP formation likely exists within repeats across many organisms, which may have been acted upon in distinct ways in different species. The combination of our analysis within and across species raises the question of whether formation of double-stranded RNA is a function for which aspects of the LINE reverse transcription machinery has been selected for. We generally support the hypothesis that repeats are selected to maintain “non-self” PAMPs, whose induction and subsequent innate sensing may act as sensors for loss of heterochromatin, avoidance of genome instability^22, 23^, or aberrant RNA processing^22, 50, 51, 53^.

While a species may have evolved to maintain PAMPs, it can be difficult to establish whether PRR signaling is the primary reason for why a PAMP evolved in the first place. Inverted Alus can be hotspots for RNA-editing, altering gene expression over evolutionary times scales, while simultaneously acting a PAMP for PRRs such as MDA5^57–59^. HSATII may have a DNA regulatory function as well, as its DNA sequences can sequester chromatin regulatory proteins and trigger epigenetic change^60^. Yet in cancer, where Alus and HSATII are often overexpressed, the same features can be sensed as PAMPs^10, 61^. Moreover, such functions are not mutually exclusive. The high CpG presence in L1 may have evolved both to allow active L1 species to remain silenced and to serve as a danger signal when aberrant demethylation occurs. For multicellular organisms with a high degree of epigenetic regulation and chromosomal organization, a repeat species with a non-immune function may be co-opted when it offers an opportunity to maintain stimulatory features to release a danger signal when epigenetic control is lost, such as during the release of repeats after p53 mutations, where immunostimulatory repeats may offer a back-up for p53 functions such as senescence^6, 62^.

Our work has several implications for how to quantify self versus non-self discrimination by the innate immune system. While we focus on motif usage and the formation of long double-stranded RNA structures, our framework is generalizable to other, more complex patterns and machine-learning approaches. Mathematically, our work highlights how approaches from statistical physics, such as maximum entropy and transfer matrix calculations can be used in efficient genome wide calculations and comparisons. The selective forces are intrinsic quantities which can be compared from sequence to sequence. Therefore, they are ideal for evolutionary analysis of genome features, the complexity of which can be added in future models. For instance, Y RNAs, implicated in RIG-I sensing during RNA virus infection^4^, have a more complicated feature set which includes an RNA modification^63^. The potential association of high CpG content with replication competent L1 may also serve as a marker for STING-cGAS^20^ activation during reverse transcription or sensing of the ribonucleoprotein complex by TRIM5α^64^, and has been implicated here as a co-sensor with *ZCCHC3*^44^. Using such methods to “decipher” noncoding genome regions and to assign them a function may allow such regions to be further exploited therapeutically. The implication is that we can learn a “repeat code” of self-agonists within our genome held by selection to stimulate receptors under specific circumstances. Such work will be enabled by emerging sequencing technologies, such as telomere-to-telomere^65^ sequencing, and broad sequencing of receptor ligand pairs. In doing so, we may discover a new set of phenotypes hiding in the non-coding genome.

## Data and Code Availability Statement

Original data will be made public upon acceptance. Code will likewise be deposited on GitHub.

## Acknowledgements

This research was funded in part through the NIH/NCI Cancer Center Support Grant P30 CA008748 (A.S., S.S., B.G.); NIH grants R01AI081848 (N.V., B.G.), R01CA240924 (A.S., B.G.), R01GM126170 (J.L.), R01AG078925 (J.L.), P50 254838-01 (O.A.-W.). and U01CA228963 (A.S., S.S., B.G.); Fondation de la Recherche Médicale: ANR-Flash Covid, Project SARS-Cov-2immunRNAs (S.C., R.M.); the V Foundation for Cancer Research (A.S.); the Mark Foundation ASPIRE award (B.G.); the Pershing Square Sohn Prize-Mark Foundation Fellowship (A.S., O.A.-W., N.V., B.G.); the Edward P. Evans Foundation (O.A-W.); the Canadian Institute of Health Research (CIHR), New Investigator salary award 201512MSH360794-228629 (D.D.C.); Canada Research Chair (D.D.C.); CIHR Foundation Grant FDN 148430 (D.D.C.); CIHR Project Grant PJT 165986 (D.D.C.); NSERC 489073 (D.D.C); and the European Union’s Horizon 2020 research and innovation programme under the Marie Sklodowska-Curie grant agreement No 101026293 (A.D.G.). The authors would like to acknowledge productive conversations with Jef Boeke, Kathleen Burns, Katherine Chiappinelli, Arnold Levine, Phil Sharp, Martin Taylor, David T. Ting, and the De Carvalho, Cocco, Greenbaum and Monasson laboratories; and thank Nicole Rusk for reading and editing the manuscript. We would also like to acknowledge support from the National Center for Dynamic Interactome Research (NIH P41GM10982); the Genome Technology Center at NYULH, a shared resource partially supported by the Cancer Center Support Grant P30CA016087 at the Laura and Isaac Perlmutter Cancer Center; and the UMCG Research Sequencing Facility and Utrecht Sequencing Facility (USEQ; USEQ is subsidized by the University Medical Center Utrecht and The Netherlands X-omics Initiative [NWO project 184.034.019]).

## Author Contributions

Conceptualization: P.S., R.M., S.C., B.G.; Research Plan: P.S., R.M., S.C., B.G; Mathematical Modeling: P.S., A.D.G.; R.M., S.C., B.G.; Double-stranded Force Calculation: P.S., A.D.G.; R.M., S.C.; Model Implementation: P.S., A.D.G.; Comparitive Genomic Analysis: A.S.; Data Analysis: P.S., A.D.G., A.S., H.L., S.M., S.S.; L1 experimental design and execution: J.L., H.J., B.H.L; Double-stranded RNA experimental design; D.D.C.; Interpretation: P.S., N.V., J.L., O.A.-W., D.D.C., R.M., S.C., B.G.; Writing: P.S., A.D.G.; R.M., S.C., B.G.; Reviewing & Editing: P.S., A.D.G., J.L., O.A.-W., D.D.C., R.M., S.C., B.G..

---

**Extended Data Figure 1.**
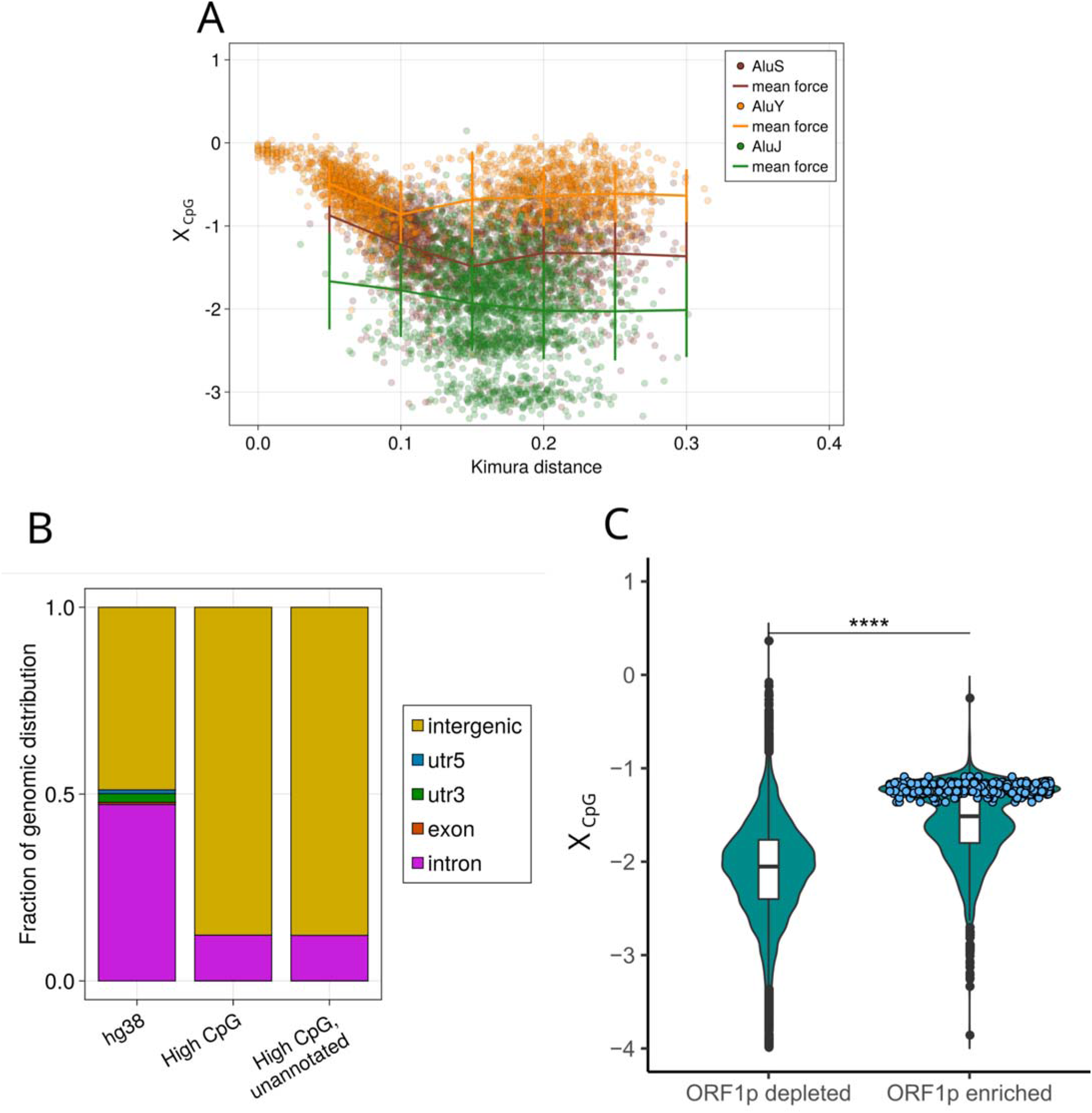
**A,** *x*_CpG_ versus Kimura distance from consensus sequence for each Alu family. Solid lines indicate binned means and standard deviations. **B,** Genomic distribution of high-*x*_CpG_ (*x*_CpG_ > 0) regions in the human genome (center), compared with the distribution of the full genome (left bar). In the right bar we show the genomic distribution of high-*x*_CpG_ regions that do not overlap with any repeat in the DFAM database. **C,** *x*_CpG_ on L1 ORF1p binding LINE transcripts in N2102Ep. Functional intact LINEs are colored in blue. BH corrected p-value is labeled for t-test. **** denote adjusted p-value < 0.01. L1 ORF1p enriched and depleted transcripts are selected by differential expression analysis between L1 ORF1p-IP vs Mock/total with a |log2FC| greater than 3 and adjusted p-value < 0.05.

**Extended Data Figure 2.**
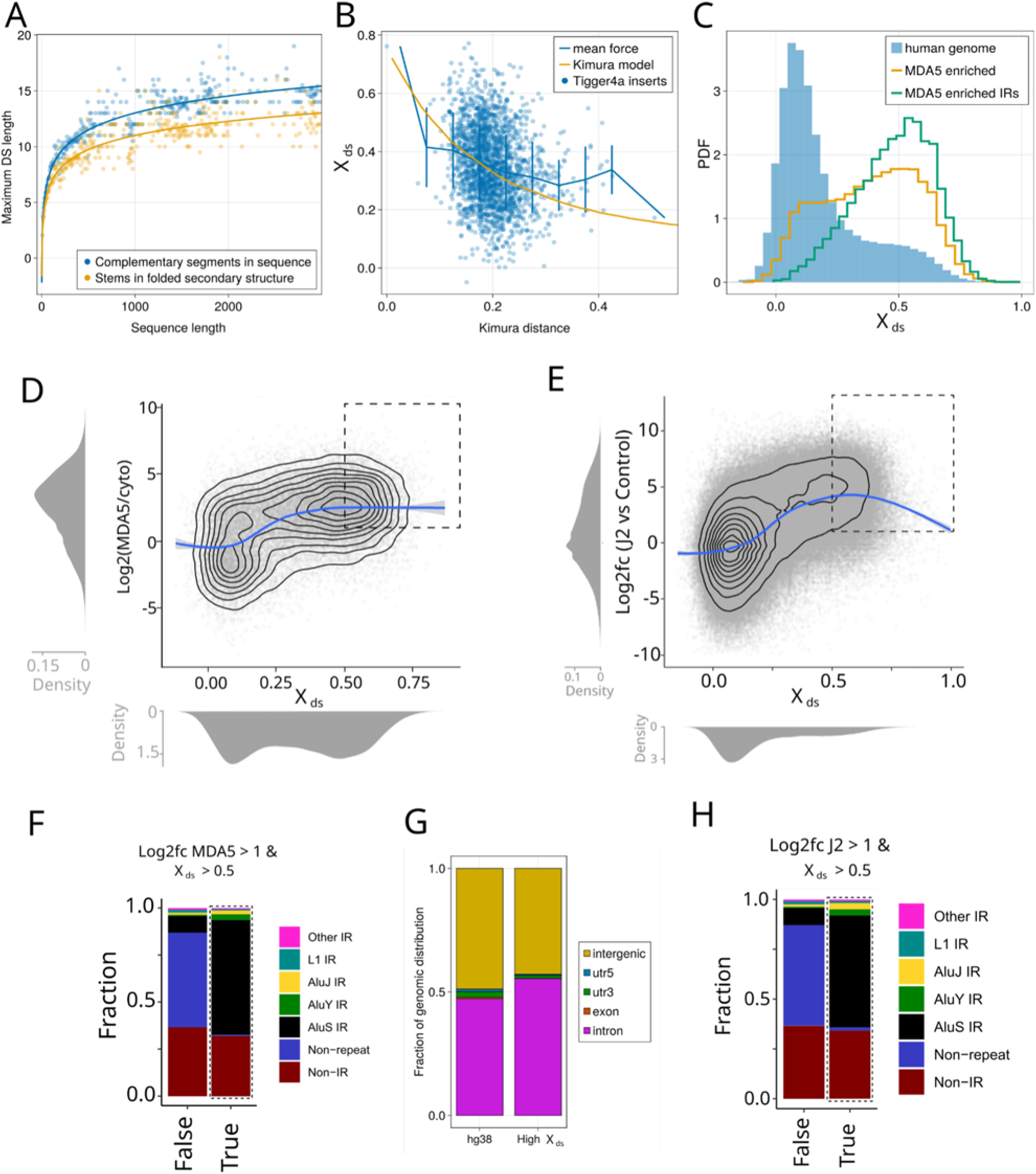
**A,** The mean of maximum lengths in a secondary structure in a single-stranded RNA sequence (green line), and the mean maximum length of complementary segments (blue line), along with respective fits (Methods). **B,** *x*_ds_ on repeat family Tigger4a. The force relaxation evolutionary model fit shows the relaxation of the inserts compared to the relaxation simulated by neutral Kimura model. **C,** *x*_ds_ histograms in human genome (sliding window with transcript of length of 3 kb) compared to MDA5 binding RNA transcripts as experimentally found in^52^. Enriched transcripts have a positive log-enrichment with respect to the control experiment. Inverted repeat (IR) transcripts are annotated repeats with another repeat of the same family in opposite genomic sense within 3 kb. **D,** Correlation between log-enrichment of reads aligning to each complementary sequence in MDA5- binding experiment, and *x*_ds_. The blue line shows the fit of a generalized additive model. **E,** relation between log-enrichment of reads aligning to each complementary sequence in J2-binding experiment, and *x*_ds_. The blue line shows the fit of a generalized additive model. **F,** Type of repeat with the longest overlapping sequence in complementary sequences with high MDA5 signal and high *x*_ds_ (*x*_ds_ > 0.5) and complementary sequences with low MDA5 signal and low *x*_ds_. **G,** Genomic distribution of high-*x*_ds_ regions in the human genome (right bar), compared with the distribution of the full genome (left bar). **H,** Type of repeat with the longest overlapping sequence in complementary sequences with high J2 signal and high *x*_ds_ and complementary sequences with low J2 signal and low *x*_ds_.

**Extended Data Figure 3.**
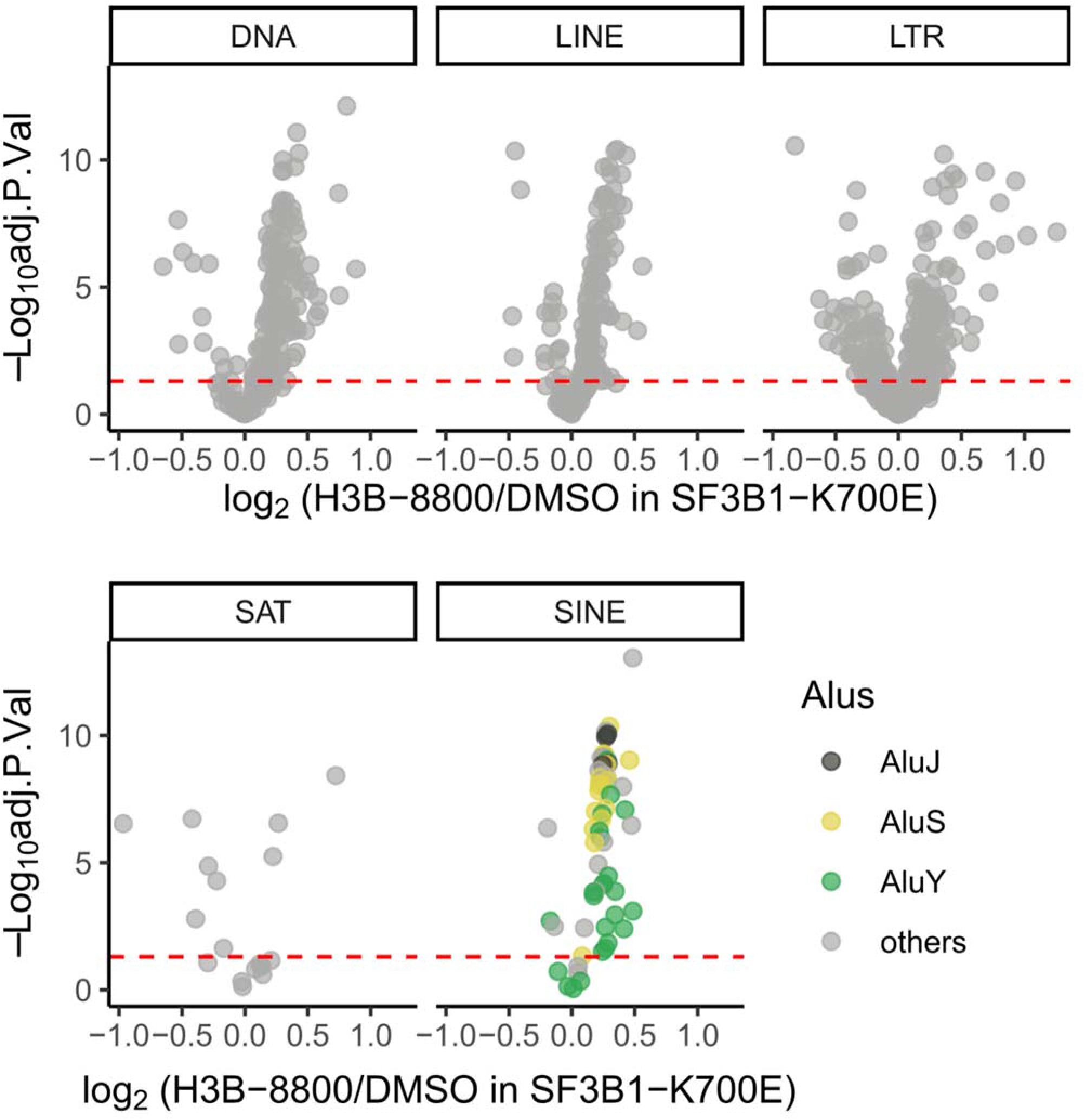
Volcano plot of repeat element expression of elements with double stranded force greater than 0.5 in H3B-8800 versus DMSO treated SF3B1-K700 mutant K562 cell lines.

**Extended Data Figure 4.**
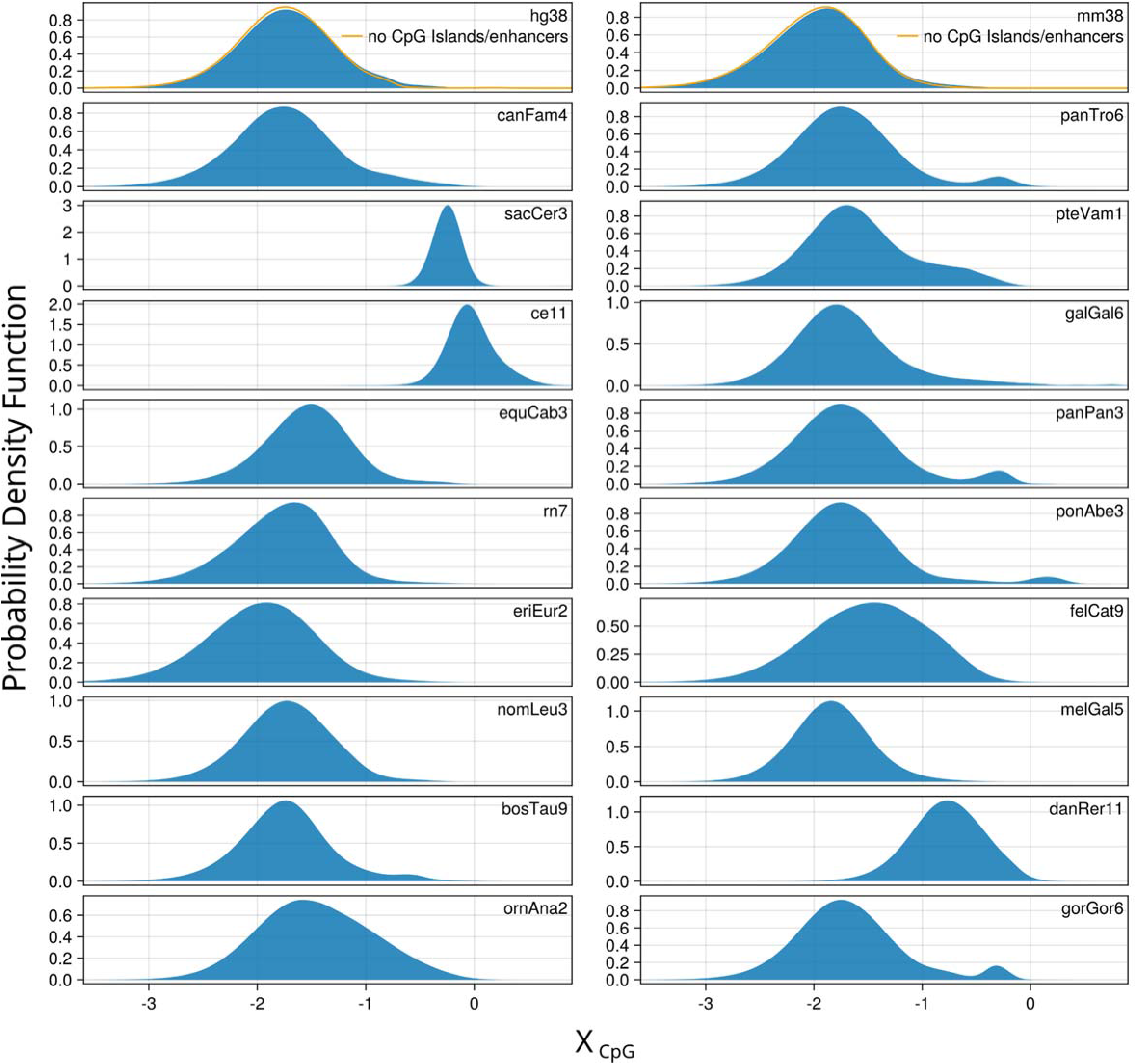
Distributions of *x*_CpG_ in several organism genomes (sliding window with transcript of length of 3000). For human and mouse, we also show, in orange, the profile of the histogram of *x*_CpG_ after excluding reads annotated as CpG islands or enhancers.

**Extended Data Figure 5.**
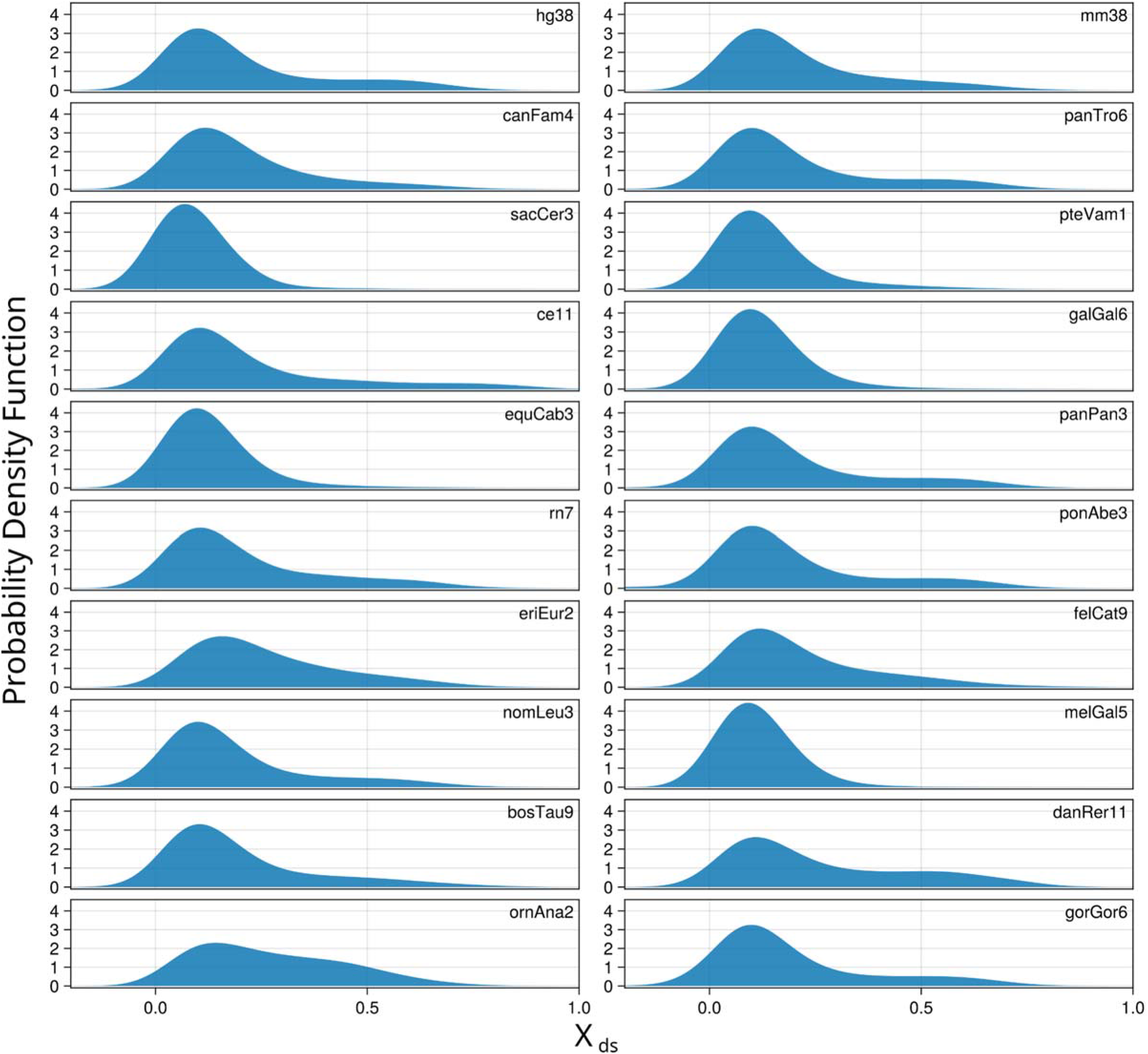
Distributions of *x*_ds_ in several organism genomes (sliding window with transcript of length of 3000).

**Extended Data Figure 6.**
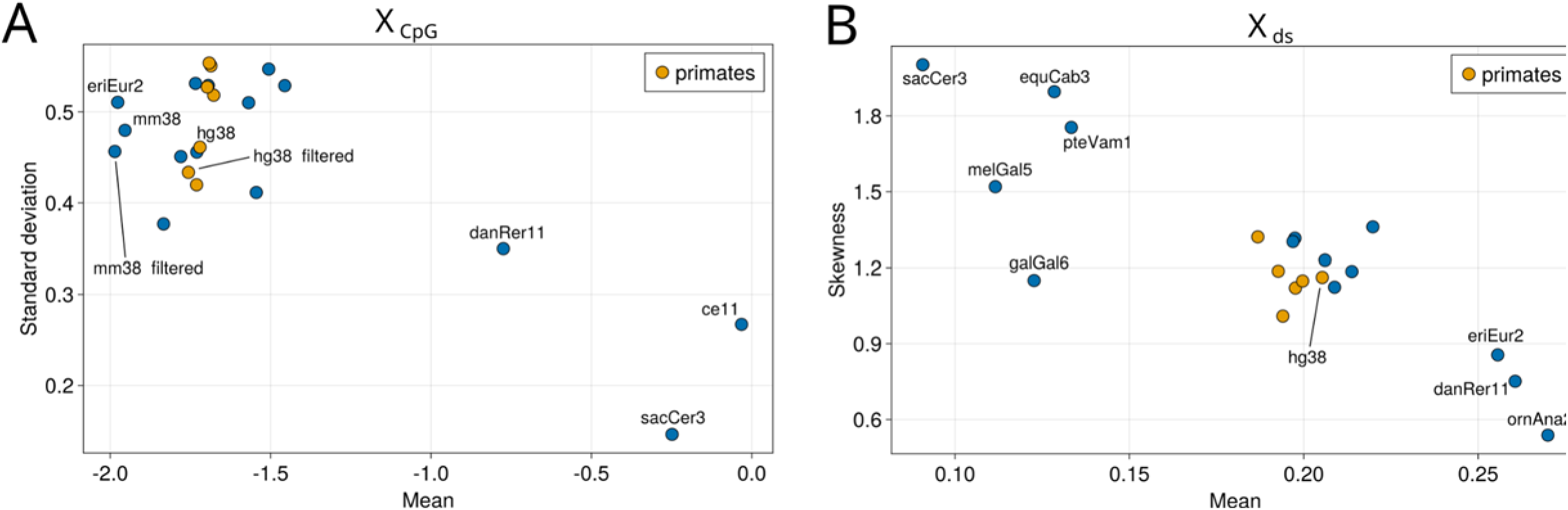
**A,** Standard deviation versus mean of *x*_CpG_ computed for each 3000-base windows for each organism analyzed. Orange denotes points computed from primate genomes. **B,** Skewness versus mean of *x*_ds_ computed for each 3000-base windows for each organism analyzed. Orange denotes points computed from primate genomes.

## Methods

### Quantification of forces on sequence features

We define a Maximum Entropy (MaxEnt) framework [1] to determine the least constrained probability distribution over the set of sequences ***s*** = *s*_1_*, s*_2_*.. . s_L_* of length *L* compatible with the observed occurrences (measurement) of a set of *M* features ***N*** = *N*_1_*, N*_2_*.. . N_m_* . The distribution is written as

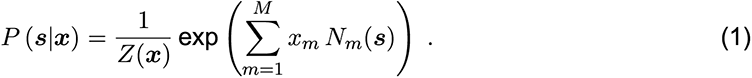

Here *Z*(***x***) is a normalization factor:

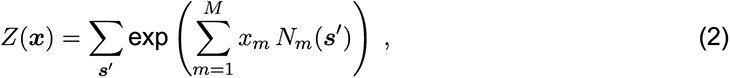

where the sum runs over all possible sequences ***s****^t^* having length *L*. The set of parameters ***x*** = *x*_1_*, x*_2_*, …, x_M_*, hereafter called *selective forces*, is chosen so that the average value of each feature over the distribution *P* matches the observed number of this feature ***N*** ^obs^ in one or more reference sequences:

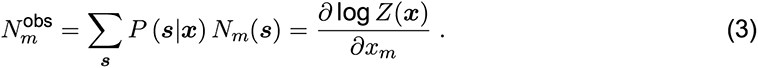

The above equalities define a set of *M* coupled, nonlinear equations, with a unique solution due to the convexity of log *Z*(***x***). The forces are analogous to chemical potentials in statistical physics.

We use our formalism to calculate selective forces on sequences due constraints imposed by: (1) the force due to potentially immunogenic CpG motifs (the CpG force, *x*_CpG_), (2) the forces on all nucleic acid motifs with length up to three nucleotides, and (3) the force on long complementary sequence stretches (the double-stranded force, *x*_ds_). For the later case we use an equivalent direct approach to simplify the calculation.

### Forces on CpG and other individual dinucleotides

To estimate the force on CpG motifs, *x*_CpG_, arising from constraints on the usage of CpG dinucleotides we derive the MaxEnt distribution *P* of sequences with five forces chosen to reproduce the frequencies *f* (*σ*) of the four nucleotides, *σ* = A,C,G,U, and the frequency of CpG dinucleotides only. Other dinucleotide forces are calculated in the same manner. We find that the single-nucleotide forces are, to a good accuracy, given by *xσ* = log *f* (*σ*) (more specifically, the forces computed in this way are in a linear relationship with those computed with a full maximum entropy model, see **Methods Fig. 1A**), which allows us to approximate the normalization factor in Eq. (2) with

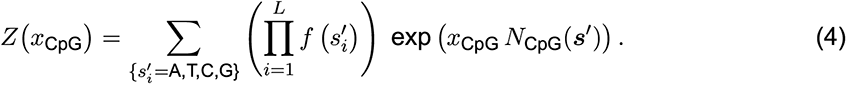

A Newton’s method based algorithm to efficiently (time *∼ O*(*L*^2^)) calculate *x*_CpG_, such that Eq. (3) (with *m* = CpG) is satisfied, was derived in [2]. A positive value of *x*_CpG_ for the observed feature count *N*_CpG_ in a given sequence indicates CpG motifs are enriched compared to what would be expected from a random sequence conditioned on single nucleotide usage only. A negative value corresponds to a depletion with respect to the null model.

### Inference of general model based on mono-, di- and trinucleotide motif usage

To infer the full set of forces on each motif of length up to 3 nucleotides we extend the formalism used for individual dinucleotide forces alone. We model each viral or repeat family as a probability distribution over the sequences characterized by the frequency of each nucleotide, dinucleotide and trinucleotide motif, as described above. In the general case, this corresponds to an overall set of *M* = 4 + 4^2^ + 4^3^ = 84 forces to infer, such as *x*_TCA_, *x*_GGC_, *x*_GA_, and *x*_T_, respectively, on the motifs TCA, GGC, GA, and T. Note that, due to symmetries of the problems, only a subset of 39 parameters can vary freely, so we are left with their inference. For instance, the sum of *N*_A_, *N*_C_, *N*_G_, and *N*_T_ is fixed (independent on the forces) and equal to *L*. However, many of these properties (such as the fact that *N*_C_ = *N*_AC_ + *N*_CC_ + *N*_GC_ + *N*_TC_) strictly hold only for infinitely long sequences, but are very well approximated in longer sequences, such as those of length equal to 3000 nucleotides frequently studied here. Because of this and similar properties, we are free to fix a certain number of forces to an arbitrary value. For instance, we can set the force of each motif containing a T to zero, without losing generality.

To infer the remaining 39 forces we used a method analogous to the one developed for CpG forces, which allows for an efficient evaluation of the maximum entropy parameters through Eqs. (2) and (3).

To train the models on viral families, we used datasets for every RNA viral family and for HIV viruses collected from the Virus pathogen Database and Analysis Resource (https://www.bv-brc.org/) [3], after removing sequences with non-standard nucleotides (different from A, C, G, T) and duplicate sequences. Influenza A viral sequences were through the Influenza Research database (now housed at [3]) and filtered with the same criteria. The model for Influenza A viruses has been trained on the sequence obtained by joining the viral segments.

The models on repeat families have been trained in the same way, using consensus repeats from [4] and grouping them in family as annotated in [5]. Each model has been trained by computing the average frequencies of each motif for the viral or repeat family, then obtaining the number of motifs for the inference procedure through multiplication by the same length for each model (5000 nucleotides).

Once the force parameters of Eq. (3) have been fitted to match the motifs statistic of a set of viral genomes or repeats in the reference list, we quantify the similarity between viral and repeat families using the symmetrized Kullback-Leibler divergence between the corresponding probability distributions *pv* and *pr* given by 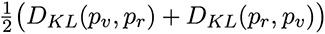 where *DKL* is the Kullback-Leibler (KL) divergence defined as

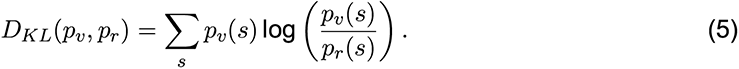

*DKL* is not an intensive quantity, as the probability distributions *pv* and *pr* depend on the length of the sequences modeled. In this work we fixed a reference length of 1000 nucleotides for all *DKL* computations.

### Computation of the Kullback-Leibler divergence

The model we define in Eq. (1) can be rewritten as

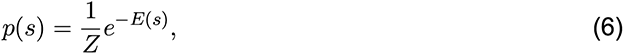

where *E*(*s*) is an energy associated to each sequence, in the usual statistical physics sense. The KL divergence can be written as

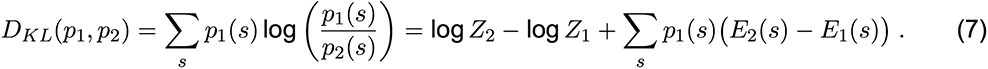

As discussed in Methods, log *Z*1 and log *Z*2 can be computed exactly with the transfer matrix method. To compute the last term on th r.h.s. of Eq. (7) we define

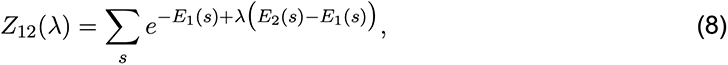

#### and we have

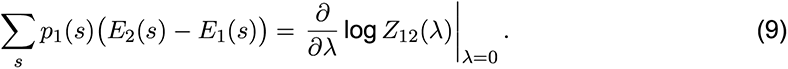

### Forces on double-stranded RNA formation

We develop a framework to quantify the length of duplex strands. Given a reference sequence ***σ*** of length *L* we compute the frequencies *f* (*σ*) of the single nucleotide *σ* = A,C,G, U. The feature *N*_ds_(***s***) is now the length of the longest subsequence of ***s*** whose complementary subsequence is also present in ***s***, and which can therefore form a duplex of the same length. We present an intuitive derivation for the forces on double-stranded RNA, *x*_ds_. That version is used preferentially in the text due to its interpretability. We also present the full MaxEnt approach, derived in an exactly parallel manner to that for forces on motifs. Due to the difficulty of computing exactly the corresponding *Z*(*x*_ds_), which further justifies the intuitive approach, we utilize an approximate calculation. We show that both approaches give directly analogous results, and can therefore be used interchangeably. Both approaches are described below, along with their formal relationship.

### Direct approach

Consider two subsequences ***s*** = (*s*1*, …, sK*) and ***s****^I^* = (*s^I^*_1_*, …, s^I^_K_*) of length *K*, with nucleotides drawn independently at random with the frequencies *f* . The probability that the two sequences are complementary is equal to *p*^compl^(*K*)= *α^K^*, with

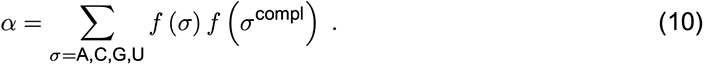

where *σ*^compl^ denotes the complementary nucleotide to *σ*. In the presence of a biasing force *x*_ds_ acting on the length of the stretch the probability that the two sequences are complementary is modified into

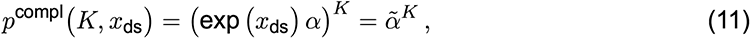

where *α̃* = exp *x*_ds_ *α*. Positive and negative forces *x*_ds_ favor, respectively, longer and shorter complementary stretches than expected from the random nucleotide null model.

Consider now a sequence of length *L*, which we partition into *N* = *L*/*K* subsequences of length *K* each. Under the simplifying assumption (that we check *a posteriori* in the following) that each pair of these subsequences is independent, the probability that none of them is fully complementary is

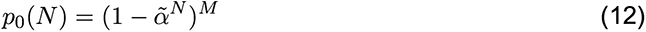

where *M* = *N* (*N* − 1)/2 is the number of pairs of segments. Equivalently, *p*0(*N*) can be interpreted as the probability that the longest fully complementary segment has length *< N* . As a consequence the probability that the longest fully complementary segment is of length equal to *N*_ds_ reads

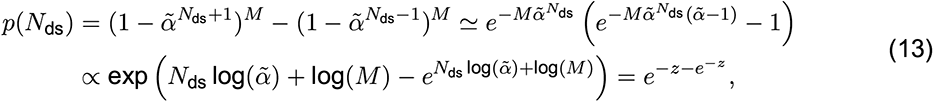

where 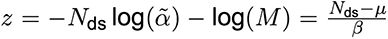 with 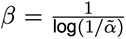 and 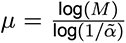. The approximations used are *α̃^N^*^ds^ 1 and *M α̃^N^*^ds^ 1, which are expected to hold in our case. Eq. (13) hence shows that the distribution of the longest fully complementary strand follows a Gumbel law, with mean *µ* + *βγ* (here *γ* is the Euler-Mascheroni constant) and variance *β*^2^*π*^2^/6.

As in Eq. (3), we can fix the force parameter *x*_ds_ by requiring that the average maximum length of fully complementary segment computed through the model, *µ* + *βγ*, is equal to the value observed in a given sequence, 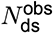, and we obtain the equation

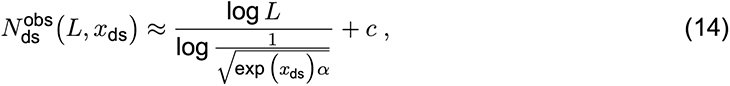

for large values of *L*. With *c* being a correction term inserted to account for the set of simplifications done, and we estimate it directly from synthetic data as follows. As a check of the validity of the expression in Eq. (14), and to estimate the value of *c*, we fit the parameters *x*_ds_ and *c* from the set of maximum length of complementary segments in randomly generated RNA sequences of lengths ranging up to *L* = 3000 bases (**Extended Data Fig. 2A**). In this work, we consider both canonical Watson-Crick pairs and Wobble pairs as complementary basepairs. We obtain *c* = − 2.2 and *x*_ds_ = 0.06, a value compatible with the zero force expected for this null model.

Eq. (14) with *c* = −2.2 can now be used to estimate *x*_ds_ for a reference sequence of length *L* (with nucleotidic frequencies *f*) and with longest complementary stretch of length 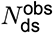. We thus obtain a single metric to compare distribution of double-stranded segments across various RNA sequence ensembles and families diverse sequence statistics and lengths.

### Maximum Entropy approach

We start from the probability that the longest fully complementary segment is of length equal to *N*_ds_, *p*(*N*_ds_). According to the maximum entropy principle, the probability distribution on sequences of length *L* which maximizes entropy while fixing the length of the maximum complementary segment is with the normalization

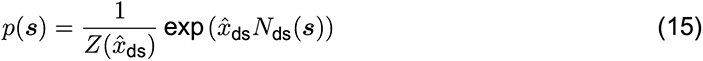

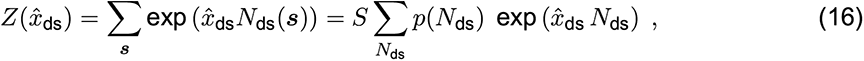

where *x̂*_ds_ is the double-stranded force acting on the length of the longest complementary stretch using the MaxEnt approach, and *S* is the total number of sequences of length *L*. Under this model, the probability of observing a sequence with maximum complementary segment of length *N*_ds_ is

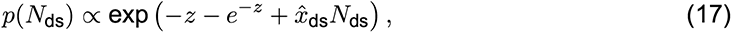

where *z* = -*N*_ds_ log(*α*) - log(*L*(*L* - 1)/2). For large *L* we can compute the average value of *N*_ds_ by integrating the continuous version of the probability distribution, and we obtain

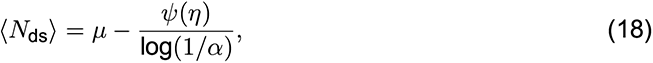

where 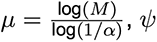 is the digamma function and *η* =1 *− βx̂*_ds_.

We can now substitute *N*_ds_ with the observed value of *N*_ds_ in the sequence under analysis, and add a constant *c′* to take care of the approximations done, and we obtain the equation

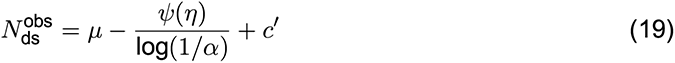

This equation can be used to estimate the MaxEnt double-strand force *x̂*_ds_. We estimated *c′* from randomly generated RNA sequences of lengths ranging up to *L* = 3000 bases and obtained the value of *−*1.79.

### Comparison of Direct and MaxEnt approaches

The direct approach to calculating *x*_ds_ is used in the manuscript. While the direct approach and the MaxEnt force are in general different, for a fixed value of *α* there is a monotonic relationship between these two quantities:

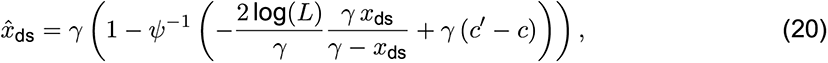

where *γ* = log(1/*α*), *c^!^* and *c* are the constants inferred for the two approaches from synthetic data, *x̂*_ds_ is the MaxEnt double-stranded force, *x*_ds_ is the double-stranded force computed via the direct approach, and *ψ^−^*^1^ is the inverse of the digamma function in the interval (0, +). In particular, we checked that even when *α* is computed for each 3000 nucleotide window, the relationship gives an extremely good approximation, thus any subsequence with double-stranded force larger than a given threshold would be equivalently characterized by high MaxEnt double-stranded force, as shown in **Methods Fig. 1B**.

**Methods Figure 1.**
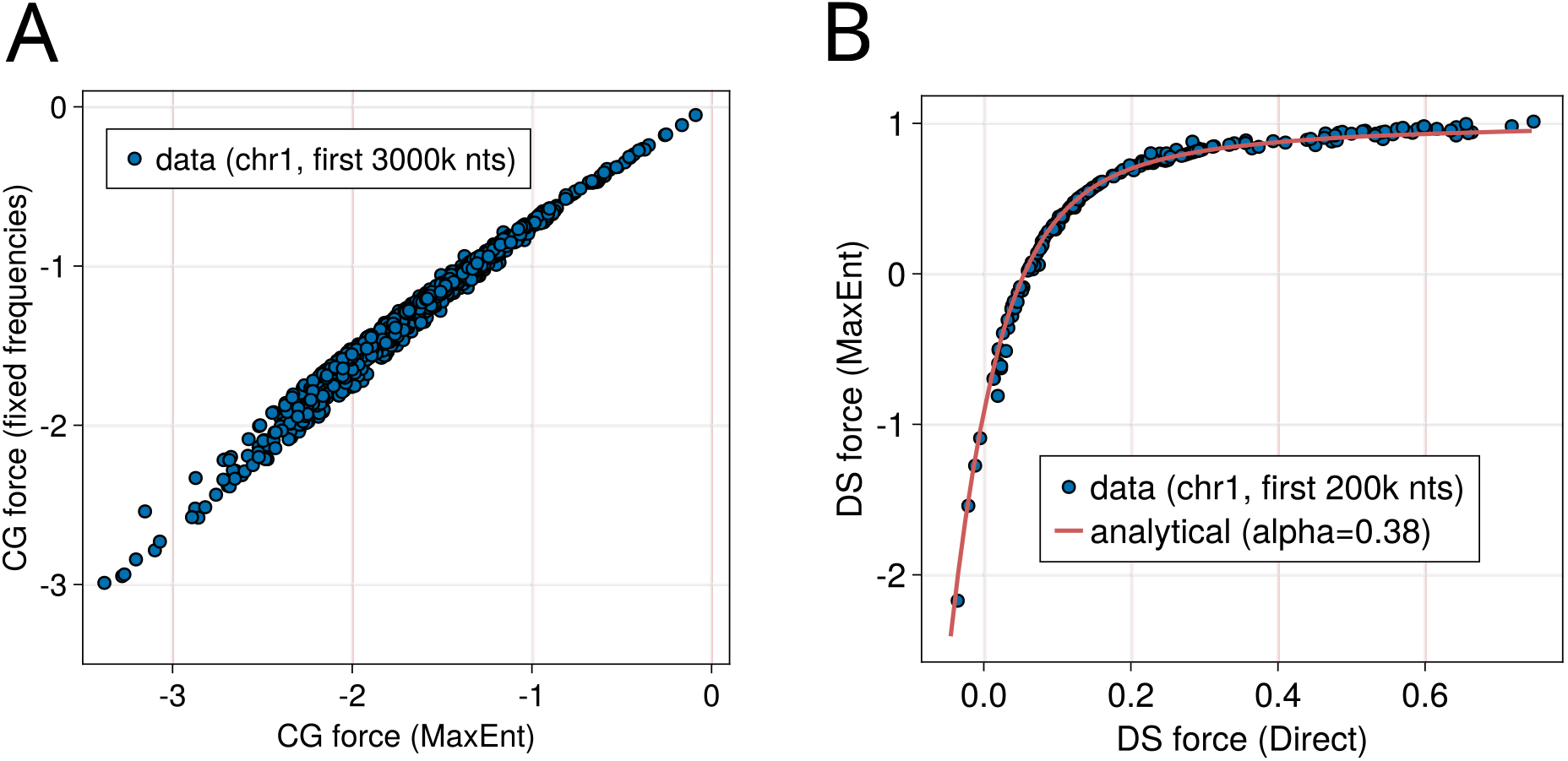
**A**, Comparison of *x*_CpG_ computed with a full-maximum-entropy model (x axis) and with the method used in this work, Eq. (4). **B**, *x*_ds_ computed with the Direct approach (horizontal axis) versus the MaxEnt approach (vertical axis), for the sliding windows within the first 200k bases of the first chromosome of the human genome. The red line correspond to the analytical relationship between the two forces computed for *α* = 0.38, which is the value obtained by considering the full human genome.

### Compressing the *x*_ds_ table

We initially computed *x*_ds_ for each sliding window of 3kb in the human genome (hg38 assembly). The resulting table of windows and associated *x*_ds_ values, however, are not appropriate for some of the successive analyses, mostly because the same pair of complementary sequences appears in many close-by windows. To deal with this issue we produced a new, compressed *x*_ds_ table with the following rules: (i) we discarded windows having the two complementary sequences less than 10 nucleotides away from the window ends; (ii) whenever more than one window have exactly the same pair of complementary sequences, we took the most upstream window and we discarded the others. Rule (i) prevents the edges of the windows from “cutting” one of the two complementary sequences, generating cluster of very similar pairs of complementary sequences in consecutive windows, while rule (ii) prevents the presence of multiple windows associated to the same pair of complementary sequences.

### Evolutionary dynamics of a sequence motif with force relaxation formalism

One can harness the formalism developed in Eqs. (1) - (3) to study an evolutionary dynamics of number of motifs *Nm*, as it approaches the steady state (equilibrium) value 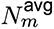 [2]. As a sequence evolves it undergoes mutations, which cause changes in the number of motifs (and hence associated value of *xs*). To model the evolutionary dynamics of sequences, we assume the number of motifs (*Nm*) evolves according to the relaxation dynamics given by

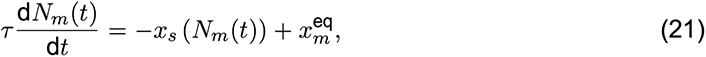

where *τ* sets the timescale. The number of motifs reaches its stationary (equilibrium) value when 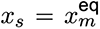, at which point the selective force is balanced by the entropic forces which randomize sequences. It is convenient to express (21) as

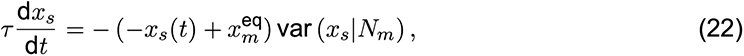

where var (*xs|Nm*) is the variance of *xs* for a given *Nm*.

If we can express var (*xs Nm*) as a function of *xs*, it is possible to obtain a solution of (22) that can then be fitted to the dataset with timescale *τ*, thus providing the approximation of relaxation dynamics, along with the estimate of the time it will take to *xs*(*t*) (and hence the number of the corresponding sequence motifs *m*) to reach its equilibrium value. For the case of HSATII and LINE-1 we fit var (*xs|Nm*) as a quadratic function of *xs*.

### Kimura-based model of population genetics for the evolution of sequence motifs

In addition to the force relaxation model introduced above, we present here a different approach to study the evolution of nucleotide sequence motifs based on the Kimura model of sequence evolution. We implement the model numerically, and evolve a set of sequences to provide a null model of neutral sequence evolution. For each simulation step, we pick a random base and mutate it to a randomly chosen different base with a given probability. We consider different possible mutation probabilities depending on the type of base it is mutating into, as well as on the context (identity of the bases in the neighborhood), as transversion (purine mutating to pyrimidine or vice versa) and transition (purine mutating to purine or pyrimidine mutating to pyrimidine) substitutions in sequences can have different likelihood [6].

Additionally, in vertebrates and plants, mutations in CpG context are known to be more common due to CpG hypermutability [7]. Hence, for the mutation rates in the model implementation, we use different ratios of mutation rates *µ*_TiCpG_:*µ*_TvCpG_:*µ*_Ti_:*µ*_Tv_ (corresponding to nucleotide transitions and transversions in CpG context and to transitions and transversion in non-CpG context). In particular, we consider the ratios introduced in Ref. [6] and which are listed in Table 1. The increased mutation rate on CpG dinucleotides has the effect of reducing the expected CpG number after relaxation of a sequence, and it can thus be related to an equilibrium CpG force. To compute this force we considered the model without the transition-transversion bias (that cannot affect the number of CpGs at equilibrium) and we can write for the number of CpG in a sequence

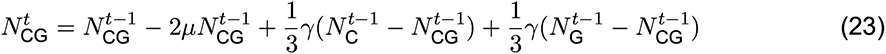

while the number of C nucleotides *NC* evolves as

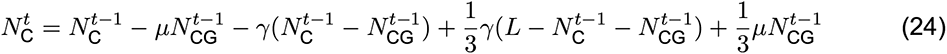

**Table 1.** Ratios of dinucleotide mutation rates (transition and transversion with and outside of CpG context) and a corresponding value of the equilibrium force on the CpG dinucleotide

| $\mu_{\text{TiCpG}}:\mu_{\text{TvCpG}}:\mu_{\text{Ti}}:\mu_{\text{Tv}}$ | $x_{\text{CpG}}^{\text{eq}}$ |
| --- | --- |
| 40:10:4:1 | -1.9 |
| 40:4:4:1 | -1.8 |
| 40:1:4:1 | -1.7 |
| 4:4:4:1 | -0.3 |
| 20:4:4:1 | -1.2 |
| 27:2:4:1 | -1.4 |

and similarly for the number *N*_G_ of G nucleotides. In these equations, *µ* is the probability of a substitution happening in a CpG context, and *γ* is the probability of the mutation happening outside a CpG context, and *L* is the length of the sequence. At equilibrium, we find

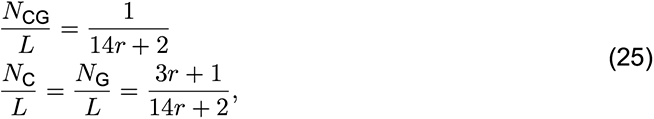

where *r* = *µ*/*γ*. We can now compute the corresponding CpG force 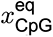 using the fact that the force is approximately equal to the logarithm of the relative frequency of the dinucleotide motif *x*_CpG_ ≈ log(*f* (CpG)/*f* (C)*f* (G)) [8]. The ratios 40:10:4:1, 40:4:4:1 and 40:1:4:1 provide the closest approximation to relaxation to the force observed in the genome. For the neutral model, we used the 40:10:4:1 ratio as it was closer to the saturated value of *x*_CpG_ of the LINE-1 elements.

### Analysis of forces across species

Values of *x*_CpG_ or *x*_ds_ were computed for each 3000 kb sliding window. Non-numeric values were excluded. Windows with one or more ambiguous characters were excluded. To compute the distribution of a force across all windows, the values of that force were sorted numerically and every 100th entry was retained (entry number 50, 150, 250, …, etc.). The distribution density was computed using a Gaussian kernel with bandwidth 0.05 as implemented in scikit-learn package [9]. The density was computed for all points within the target interval ([-5:2] for *x*_CpG_, and [-2:3] for *x*_CpG_) with a step of 0.005. We compute the FDR as the ratio of the cumulative area of the null model distribution to the right of the cutoff to the cumulative area of the distribution to the right of the cutoff. The null model distribution is fitted as a Gaussian distribution with the peak at the point with the maximal density, standard deviation was computed using the 20 points to the right of the peak. We computed these values across the species: Pan troglodytes troglodytes, Pan paniscus, Gorilla gorilla gorilla, Pongo abelii, Nomascus leucogenys, Canis lupus, Danio rerio, Mus musculus, Rattus norvegicus, Equus caballus, Bos taurus, Gallus gallus, Felis catus, Pteropus vampyrus, Caenorhabditis elegans, Saccharomyces cerevisiae, Meleagris gallopavo, Erinaceus europaeus, and Ornithorhynchus anatinus, in addition to humans.

For humans and mice we also performed an additional analysis which excluded both enhancers and CpG islands. Coordinates of enhancers from the FANTOM database were lifted from hg19 to hg38, and mm9 to mm38 using LiftOver tool from UCSC ([10, 11], [12]). Coordinates of CpG islands for hg38 and mm38 were downloaded from UCSC. ”Filtered” data for hg38 and mm38 in the CpG plot consist only of the windows which have zero overlap with CpG islands and enhancers.

### Analysis of evolutionary conserved sequences with high *x*_CpG_ or *x*_ds_

We considered the genomes of 5 species (Pan troglodytes troglodytes, Pan paniscus, Gorilla gorilla gorilla, Pongo abelii, Nomascus leucogenys) in addition to the human genome to look for conserved regions with high *x*_CpG_ or high *x*_ds_. After computing both forces for each species exactly as we did for the human genome, we considered the set of *x*_CpG_ >0 and *x*_ds_ >0.5. We reduced the number of the CpG windows by clustering together all those which overlapped more than 1000 bases, and from each cluster we only considered the window with the highest value of *x*_CpG_. For the *x*_ds_ windows, we first compressed them by excluding windows with cut-off complementary sequences or with identical complementary sequences (as discussed above), then for each window we extracted the subsequence spanning the pair of complementary sequences. Finally we clustered together all overlapping sequences and from each cluster we only considered the window with the highest value of *x*_ds_.

We then ran BLAST to compare each of these sequences with high *x*_CpG_ or high *x*_ds_ extracted from the human genome and we retained any significant match (whenever one sequence of a given organism matched with more than one human sequences we only kept the match with the highest BLAST score) [13]. The result of this procedure consists in two sets of sequences for each organism that are alignable to human sequences, one for the high *x*_CpG_ and one for the high *x*_ds_. We then computed the overlaps between the set *A* of high *x*_CpG_ (or high *x*_ds_) organism sequences and that of the human, *B*, defined as

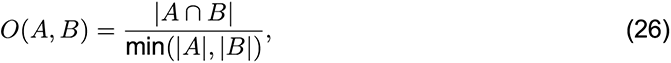

where a sequence of *A* belongs to the intersection *A ∩B* if and only if it is alignable with significant score by BLAST to a sequence of *B*. *|A|* is the size of the set *A*.

We next compared the overlaps we obtained with the time since the most recent common ancestor, taken from [14]: 6.6 million of years ago for Homo Sapiens, Pan troglodytes troglodytes and Pan paniscus; 8.9 for Homo Sapiens, Pan troglodytes troglodytes, Pan paniscus and Gorilla gorilla gorilla; 13.8 for Homo Sapiens, Pan troglodytes troglodytes, Pan paniscus, Gorilla gorilla gorilla and Pongo abelii; 16.3 for all the species considered here. Afterwards we focused on the annotations in RepeatMasker [5] and DFAM [4] databases to check for potential overlaps of the conserved sequences with annotated repeats in the human genome. For the set of high-CpG sequences we used the DFAM dataset as we found a better annotation for HSATII with respect to the one in RepeatMasker. We associated each conserved high-CpG sequence with a repeat if the corresponding window overlapped with its position as annotated in the database (for the windows overlapping with more than one annotated repeats, the one with the largest overlap was considered).

For the set of *x*_ds_ sequences we used the RepeatMasker dataset [5]. In this case, each *x*_ds_ window is characterized by two fully complementary sequences, and we searched for repeats overlapping with each of them (when overlaps with multiple repeats were found, the one with the largest overlap was considered). We observed 3 different cases: one or two of the two sequences do not overlap with any repeat annotated in RepeatMasker; each sequence overlapped with a repeat, both repeat being of the same family and annotated in the two strands of the genome, e.g. one sequence overlapping with AluS+ and one with AluS- (IR repeats); each sequence overlapped with a repeat, but of different families or of the same family but on the same strand of the genome (non IR repeats).

### Sequence ensembles

The LINE-1 sequences were obtained from L1Base2 database [15]. We separately downloaded all the sequences annotated as full-length intact and hence are more likely to still be active (146 for human genome and 2811 for mouse genome), and sequences annotated as full-length nonintact (13148 for human genome and 14076 in mouse genome). We separately aligned each of the non-intact sequences with each of the respective intact sequences using pairwise alignment and calculated the Kimura distance between the sequences [16]. We then calculated the average distance for each of the non-intact sequences from the intact-sequences, and furthermore calculated the number of CpG motifs in each sequence.

Sequences of all inserts of HSATII and all other Human Genome repetitive elements considered in this work have been obtained from the DFAM database [17] (version introduced in 2016). Each family of sequences in the DFAM database contains sequences of all its inserts in the human genome and their consensus sequence, as well as with the hidden Markov Chain Model (HMM) that we use to align inserts with respect to the consensus sequence. For comparison of sequences of inserts with respect to their consensus sequence, we only consider inserts of length longer than 150 bases. To quantify the difference between the insert sequence and the consensus sequence, we use the Kimura distance [16] between the consensus and its inserts.

We note that we use the Kimura distance [18] from the consensus sequence (for inserts from DFAM) or from average of all full-length non-intact sequences (for LINE-1s from L1Base2) as a measure of time, assuming that it is proportional to the time since insertion of the particular transposable element into the species genome. All the sequences studied in this work have been obtained from hg38 genome assembly.

### Search of long transcripts with complementary regions

We scanned the hg38 genome assembly for transcripts that can be possible source of long duplex formation. To this aim, for each window of length 3000 bases (taken in the positive sense of the read), we calculate the double-stranded force *x*_ds_ from Eq. (14), using the window-specific nucleotide frequencies to obtain *α* from Eq. (10). We considered windows resulting in *x*_ds_ *>* 0.5 ashaving a high double-stranded force when compared to the rest of the genome.

### Sequence complexity quantification

We use an approximation of Kolmogorov complexity [19] to quantify how “non-trivial” complementary segments are. Adopting the approach from Ref. [20], we use the size (in bytes) of the sequence compressed with gzip software as a proxy of the Kolmogorov complexity. Simple sequences, *e.g.* poly(AT) or poly(C) and poly(G), will have low complexity, as they can be compressed to a smaller size than a completely random sequence of the same length (which would have maximum complexity).

### Estimate of genome regions with high double stranded force

To estimate *x*_ds_ for a given repeat loci, we intersect each repeat loci with the calculated 3kb genomic windows that have high dsRNA forces (*x*_ds_ *>* 0.5). The Start and End coordinates of the corresponding dsRNA sequence pairs, which overlap with the repeat loci that match the criteria: log2FC(treated/untreated) > 0.5 and FDR < 0.05, were used to annotate different genomic features. We counted the genomic features of the predicted double-stranded RNA sequences that overlap with the upregulated repeats (log2FC > 0.5 and FDR < 0.05), and of those that overlap with the downregulated repeats (log2FC < -0.5 and FDR < 0.05). These counts have been compared with the genomic feature counts of all dsRNA sequences that overlap with the transcribed repeats to calculate the odds ratio and p-value using the Fisher Exact test.

### Transcriptome analysis

#### Analysis of repeats from splice inhibitors

Raw RNAseq data (GSE95011) associated with the Seiler, et al., 2018 study [21] were downloaded from NCBI. Briefly, reads were trimmed and quality checked using skewer first and then mapped to the human genome (hg38) and repetitive elements from RepBase [22, 23]. In quality check, Illumina reads were trimmed to remove N’s and bases with quality less than 20. After that, the quality scores of the remaining bases were sorted, and the quality at the 20th percentile was computed. Reads quality less than 15 at the 20th percentile or shorter than 40 bases were discarded. Only paired reads that passed the filtering step were retained. Only paired reads which both pass the quality check were mapped to the reference genome (hg38) using STAR (v2.7) with default parameters. Gene counts were assigned based on Gencode annotation using featureCounts (Subread package) with the external Ensembl annotation. Repeats counts per element subfamily were primarily quantified against RepeatMasker using featureCounts and then adding the counts of the unassigned reads that mapped to Repbase consensus sequence. Repeat counts of a given family is the sum of mapped reads to RepeatMasker and unmapped reads against Repbase.

#### Counts filtering, normalization and statistical analysis

Gene expression in terms of log2-CPM (counts per million reads) was computed and normalized across samples using the TMM (trimmed-mean of M-values) method using the calcNormFactors() and cpm() functions from edgeR package [24]. These low-count values (CPM < 2) were removed before calculating the size factor for each sample. Then, filtered CPM was log2 transformed and used in heat-map visualization and downstream statistical analysis. On the heatmap, genes (rows) were scaled by z-score scaling. Heat maps were generated by the R statistical programming package. Differential expression analysis was performed using limma package [25] between splicing modulator H3B-8800 treated versus DMSO treated SF3B1-K700 mutated cell line k562 for a given locus. The adjusted p-values were calculated using the Benjamini-Hochberg correction [26].

#### Analysis of MDA5 binding transcripts

We have used the double-stranded force calculation to score RNA-Seq transcripts identified in Ref. [27] to bind to MDA5 receptors. We have further looked if any of the identified transcripts that bind MDA5 from Ref. [27] are also overlapping with transcripts that have been identified as MDA5 ligands in Ref. [28].

### RNA extractions and co-immunoprecipitations (RIP) from EC cells

Embryonal carcinoma cells were cultured at 37° C in humidified incubators maintained with 7% CO2 atmosphere. N2102Ep Clone 2/A6 cells (Merk, #06011803) were cultured in DMEM (high glucose, no sodium pyruvate; Thermo Fisher, #11965092), supplemented with 10% (v/v) fetal bovine serum, 1x penicillin/streptavidin and 2 mM Glutamine (Thermo Fisher, #25030024). Large-scale growth, harvesting, cryo-milling, and co-IP was achieved as previously described [29–31], summarized as follows. α-ORF1p-, α-ORF2p-, and α-ZCCHC3-targeted co-IPs used in-house made [32] magnetic affinity media: for α-ORF1p [15 µg antibody / mg magnetic beads], we used the 4H1 monoclonal antibody (Millipore Sigma, #MABC1152); for α-ORF2p [10 µg / mg magnetic beads] we used the clone 9 monoclonal antibody [33]; for α-ZCCHC3 [10 µg / mg magnetic beads] we used the rabbit polyclonal antibody (Proteintech, #29399-1-AP). Co-IPs were conducted using 100 mg cell powder, extracted at 25% (w/v) in 20 mM HEPES pH 7.4, 500 mM or 300 mM NaCl, 1% (v/v) Triton X-100, 1x protease inhibitors (Roche, #1187358001), and 0.4% (v/v) RNasin (Promega,

#2515). Centrifugally clarified cell extracts were incubated with affinity medium (20 µl of slurry for α- ORF1p and α-ORF2p, and 15 µl of slurry for α-ZCCHC3) for 30 minutes at 4° C. The solutions were made with nuclease-free H2O and experiments were conducted using nuclease-free tubes and pipette tips. Macromolecule extractions performed on this cell line as described typically yielded between 450 - 500 µl of soluble extract at 6 - 8 mg/ml of protein as assessed by Bradford assay (Thermo Fisher, #23200). After target capture, washing the media was performed with the same solution without protease inhibitors and with RNasin at 0.1% (v/v). RNAs were eluted from the affinity media after RIP with 250 µl of TRIzol Reagent (Thermo Fisher, #15596026). After adding chloroform to the TRIzol eluate, the separated aqueous phase (containing RNAs) was obtained using Phasemaker tubes (per manufacturer’s instructions; Thermo Fisher, #A33248), and was then combined with an equal volume of ethanol and further purified using a spin-column according to the manufacturer’s instructions (Zymo Research, #R2060). For α-ZCCHC3 co-IP, two 100 mg-scale preparations were pooled prior to spin column purification. Eluates from α-ORF1p and α- ORF2p co-IPs were not treated with DNase I on-column, this was done during the sequencing library preparation (described, below); eluates from α-ZCCHC3 were DNase I treated on-column. Purified nucleic acids from α-ORF1p and α-ORF2p co-IPs were eluted in 6ul of nuclease-free water; purified nucleic acids from α-ZCCHC3 IPs were eluted in 10ul of RNase-Free water; in all cases 1 µl was used for quality analysis and the remainder conserved for RNA-seq. Mock RNA co-IP controls were prepared in an identical manner using either naïve polyclonal mouse IgG (control for α-ORF1p; Millipore Sigma, #I5381) or naïve polyclonal rabbit IgG (control for α-ORF2p: Innovative Research, #IRBIGGAP10MG; control for α-ZCCHC3: Millipore Sigma, #I5006). Total RNA controls were prepared by combining up to 35 µl of the clarified cell extracts with up to 500 µl of Trizol, vortex mixing for 1 min, then snap freezing in liquid N2 - and then later proceeding as above.

#### cDNA library preparation and RNA-seq

All the sequenced samples/replicates that are reported in this study are listed in Supplementary Table 6.

#### α-ORF1p and α-ORF2p RIP-seq

RNA extractions were quantified and quality controlled using RNA Pico Chips (Cat. #5067-1513) on an Agilent 2100 BioAnalyzer. RNA-Seq cDNA libraries were prepared using the Trio RNA-Seq

Library Prep kit (Tecan, #0357-A01) with AnyDeplete Probe Mix-Human rRNA (Tecan, #S02305). DNase treatment preceded cDNA synthesis. cDNA synthesis: 3 - 5ng of input RNA from α-ORF1p RIPs and mock IPs, 1 ng from α-ORF2p RIPs and mock IPs, and 50 ng of total RNA were used with 8 (2+6) cycles of pre-depletion PCR library amplification and 8 (2+6) cycles of post-depletion amplification; the libraries were purified using Agencourt AMPure XP beads (Beckmann Coulter), quantified by qPCR, and the size distribution was checked using the Agilent TapeStation 2200 system. Final libraries were sequenced, paired-end, at 50 bp read-length on an Illumina NovaSeq 6000 v1.5 with 2% PhiX spike-in.

#### α-ZCCHC3 RIP-seq

RNA extractions were quantified and quality controlled using an Agilent TapeStation 4200 and High Sensitivity RNA ScreenTape (Agilent, #5067-5579). RNA-seq cDNA libraries were prepared using the SMARTer Stranded Total RNA-Seq Kit v3 - Pico Input Mammalian (Takara, #634485), including rRNA depletion during library construction. cDNA synthesis: 5 ng of input RNA from α-ZCCHC3 RIPs and total RNA, and 1 ng of input RNA from mock IPs were used with 5 cycles of pre-depletion PCR amplification and 12 cycles (α-ZCCHC3 RIPs and total RNA) or 14 cycles (mock IPs) of post-depletion amplification. Libraries were purified using NucleoMag beads supplied in the library preparation kit and subsequently quantified using the Qubit 4 Fluorometer and the Qubit dsDNA HS assay kit (Invitrogen, #Q32854). The size distribution was checked using the TapeStation 4200; noting that primer dimers (~150bp) were persisted in the mock IP libraries (motivating an additional round of cleaning). To treat all libraries equally, they were pooled in a 4:4:1 ratio (α- ZCCHC3 RIPs:total RNA:mock IP) based on molarity of fragments of interest (range 200-1000 bp). An additional round of cleanup with NucleoMag beads was done to remove the primer dimers. A size selection of the final library pool was performed on a 2% E-gel EX (Invitrogen, #G401002) to exclude small fragments (less than about 200bp) and the DNA was eluted from the gel slices using the Zymoclean Gel DNA Recovery Kit (Zymo, #D4001), followed by quantification (Qubit) and quality control (TapeStation). Final libraries were sequenced, paired-end, at 250bp read-length on an Illumina NovaSeq 6000 platform.

#### RIP-seq read mapping and quantification

Reads were trimmed and quality checked using skewer [34]. Briefly, ends of the reads were trimmed to remove Ns and bases with quality less than 20. After that, the quality scores of the remaining bases were sorted, and the quality at the 20th percentile was computed. Reads were discarded if their quality at the 20th percentile was less than 15. In addition, reads shorter than 40 bases after trimming were discarded. If at least 1 of the reads in the pair failed the quality check and had to be discarded, we discarded the mate as well. Quality filtered reads were mapped to annotated repeat loci in RepeatMasker using software: Quantifying Interspersed Repeat Expression (SQuIRE) (https://github.com/wyang17/SQuIRE) [35]. Briefly, the SQuIRE pipeline first obtains reference annotation files from RepeatMasker, then aligns reads using STAT and, lastly, quantify locus-specific repeat expression by redistributing multi-mapping read fractions in proportion to estimated TE expression with an expectation-maximization algorithm.

#### CpG quantification for repeats in hg38

Sequences of repeats were extracted from hg38 based on the coordinate annotation in Repeat- Masker. *x*_CpG_ was then calculated for those hg38 derived repeat sequences. L1 inserts reported in RepeatMasker are considered intact whenever they have minimum 80% overlap with inserts annotated as full-lengt intact in the L1Base database.

#### Selection of RIP-Seq enriched repeats/transcripts

Samples extracted at 300mM and 500mM NaCl were used as replicates to increase statistical power after checking for transcript composition similarity (via hierarchical clustering). Targeted protein enriched transcripts were selected by Log2(Co-IP/mock) > 3, Log2(Co-IP/total RNA) > 3, and Benjamini-Hochberg adjusted p-value < 0.05. Similarly, target protein depleted transcripts were selected by Log2(Co-IP/mock) < -3, Log2FC(Co-IP/total RNA) < -3, and Benjamini-Hochberg adjusted pvalue < 0.05.

### Immunoprecipitation of double-stranded RNA by J2 antibody

#### Experimental protocol

Protein G Dynabeads were washed twice and resuspended in antibody conjugation buffer (1x PBS, 2mM EDTA, 0.1% BSA (w/v). 5 µg of anti-dsRNA mAb (J2) (SCICONS, cat# 10010500) were bound to 30 µl of washed beads overnight at 4° C on a rotating wheel. 10^7^ Patient-derived POP92 cells per IP were fixed with 0.1% paraformaldehyde at room temperature (RT) for 10 min. Immediately, cells were quenched by adding glycine and washed twice with cold PBS. Crosslinked cells were lysed in lysis buffer (20mM Tris [pH 7.5], 150mM NaCl, 10mM EDTA, 10% Glycerol, 0.1% NP-40, 0.5% Triton X-100, supplemented with protease inhibitor tablet) for 15 minutes on ice. Following a spin at 12,000*g* at 4° C for 15 minutes, supernatant was transferred to a new eppendorf. The lysate was then immunoprecipitated using 30 µL antibody-conjugated Dynabeads per IP reaction overnight at 4° C in a rotator. Following magnetic separation, beads were washed three times with high salt wash buffer (20 mM Tris pH 7.5, 500 mM NaCl, 10 mM EDTA, 10% glycerol) and resuspended in 1X TBS. Per IP, 2 µL of Promega RNasinPlus RNase inhibitor (Fisher Scientific, PRN2611) and 0.5 µL of proteinase K (NEB, P8107S) was added. Decrosslinking was performed for all the IP samples at 65° C for 15 minutes. The Direct-zol RNA MiniPrep kit (Zymo Research, R2051) was used to extract RNA from IP supernatant. Samples were treated with Turbo DNase to remove any DNA contamination in the extracted RNA. Library prep was performed using Illumina Stranded total RNA ligation with Ribo Zero plus according to the manufacturers protocol. Samples were sequenced on a NovaSeq 6000 using paired end reads with 100 cycles.

#### Analysis of J2 immunoprecipitation RNAseq

RNAseq controls not enriched with J2 antibody from untreated POP92 cells were downloaded from GEO (submission GSE145639, samples: GSM4322694 and GSM4322693) [27]. 25 bp of J2 RNAseq reads were cut off with cutadapt to match the length of RNAseq control reads. All samples were aligned to the human genome hg38 using STAR with default settings [36]. BAM files were sorted using samtools. The compressed *x*_ds_ table was used to count fragments in the RNAseq data, see above for details on how this table was generated. The table describes windows for the genome as well as a complementary sequence which can form a double stranded sequence. Every complementary sequence has a seqA and a seqB part which was split into two different files. featureCounts was used to count the number of fragments aligning to seqA and seqB in J2 and RNAseq control BAM files including information about strand and reporting multimapping and muti-overlapping reads as fractional counts [37]. Counts for each seqA and seqB were then merged for each complementary sequence. For each complementary sequence, log2FC (mean J2 / mean control) was calculated and plotted against *x*_ds_ using geom_point, geom_density_2d and geom_smooth from the R package ggplot2 [38].

#### Analysis of MDA5 protection assays

Raw sequencing data was downloaded from GSE103539 and GSE145639 [28] and aligned to hg38 using STAR. The following settings were used to increase the mapping due to the repetitive nature of the data [27]: –outFilterMultimapNmax 1000 –outSAMmultNmax 1 –outFilterMismatchNmax 10 –outMultimapperOrder Random –winAnchorMultimapNmax 1000. After mapping, the data were processed as described above for J2 immunoprecipitation.

## Supplementary Material

### Supplementary Tables

Supplementary Tables are available separately in Comma Separated Value (csv) format. The description of the respective tables is provided below:

- Supplementary Table 1 contains the *x*_CpG_ and *x*_ds_ values calculated for each repeat family annotated in the DFAM database. For each family, it includes the forces of the consensus sequence and the mean calculated for the inserts in the human genome.
- Supplementary Table 2 contains the list of high-*x*_CpG_ (*x*_CpG_ *>* 0) windows of 3 kb detected in the human genome, after filtering so that windows do not overlap more than 1 kb. The table also includes information about the repeat that maximally overlaps with each window as annotated in DFAM. Finally we report the most-recent common ancestor time of the primates for which we observed high-*x*_CpG_ sequences alignable by BLAST with each high-*x*_CpG_ window in the human genome.
- Supplementary Table 3 contains the list of the unique double-stranded segments resulting in windows with high-*x*_ds_ (*x*_ds_ *>* 0.5) detected in the human genome. The processing of data is described in Methods. The table also includes information about the repeats that maximally overlaps with each of the two double-strand forming segments as annotated in RepeatMasker. Finally we report the most-recent common ancestor time of the primates for which we observed pair of segments resulting in high-*x*_ds_ and alignable by BLAST with each pair of high-*x*_ds_ segments in the human genome.
- Supplementary Table 4 contains the list of the unique double-stranded segments resulting in windows with high-*x*_ds_ (*x*_ds_ *>* 0.5) detected in the zebrafish genome (danRer11).
- Supplementary Table 5 contains the results of differential expression analysis on all repetitive elements between treated versus untreated SF3B1 mutant samples.
- Supplementary Table 6 contains the list of annotated samples subject to L1 ORF1p/ORF2p RIP-seq analysis. Salt concentration, cell type, antibody type, replicate and sequencing details are listed.

### Supplementary Figures

**Supplementary Figure S1:**
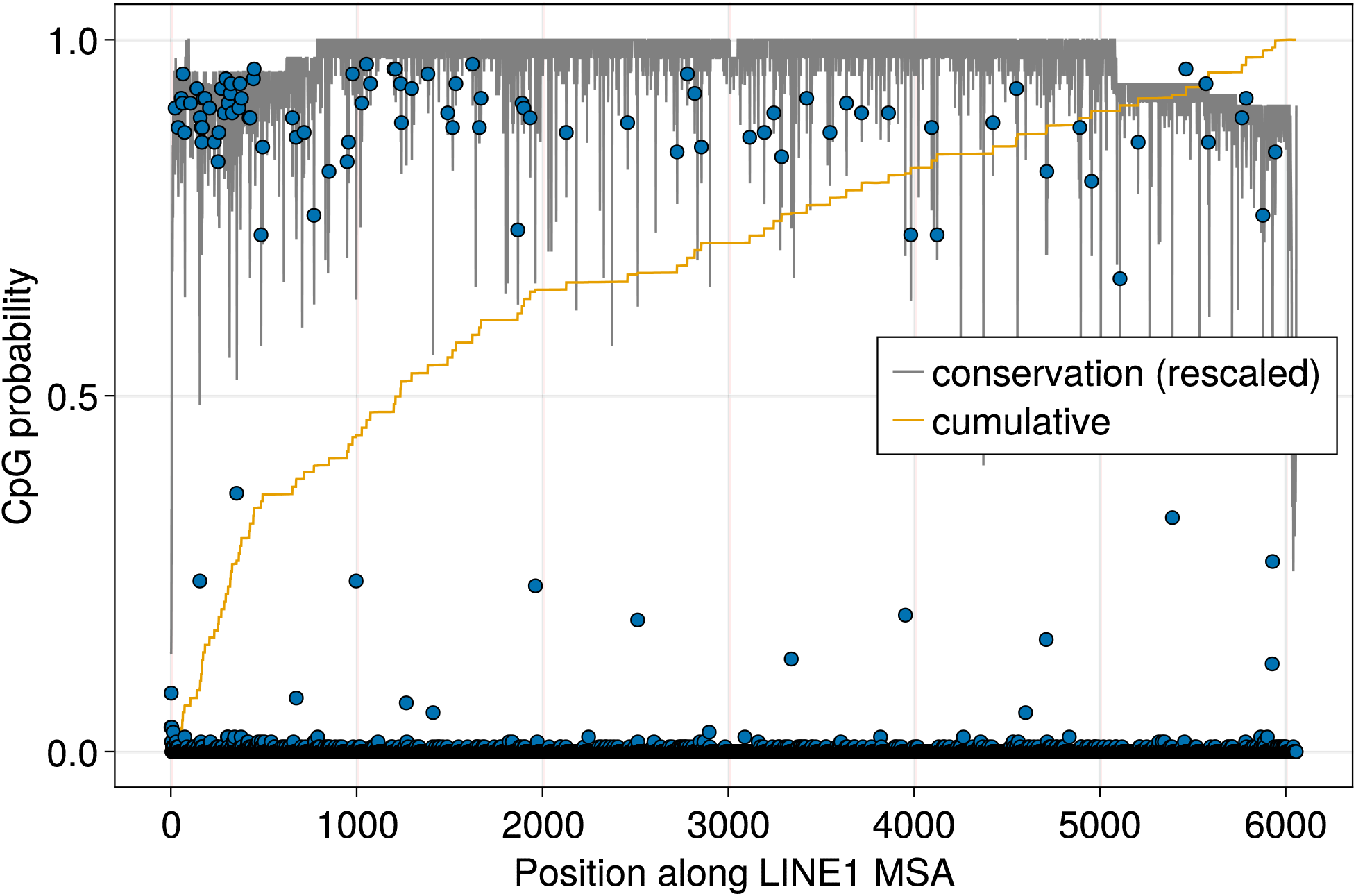
CpG occurrence probability in a multiple sequence alignment of full-length intact LINE1 inserts in the human genome. The orange line denotes the cumulative probability, and the gray line denotes the conservation (in bits) rescaled between 1 and 0: 1 + *_σ_ fi*(*σ*) log_2_(*fi*(*σ*))/ log(5), where *fi*(*σ*) is the fraction of times the nucleotide *σ* (or a gap) appears in position *i*.

**Supplementary Figure S2:**
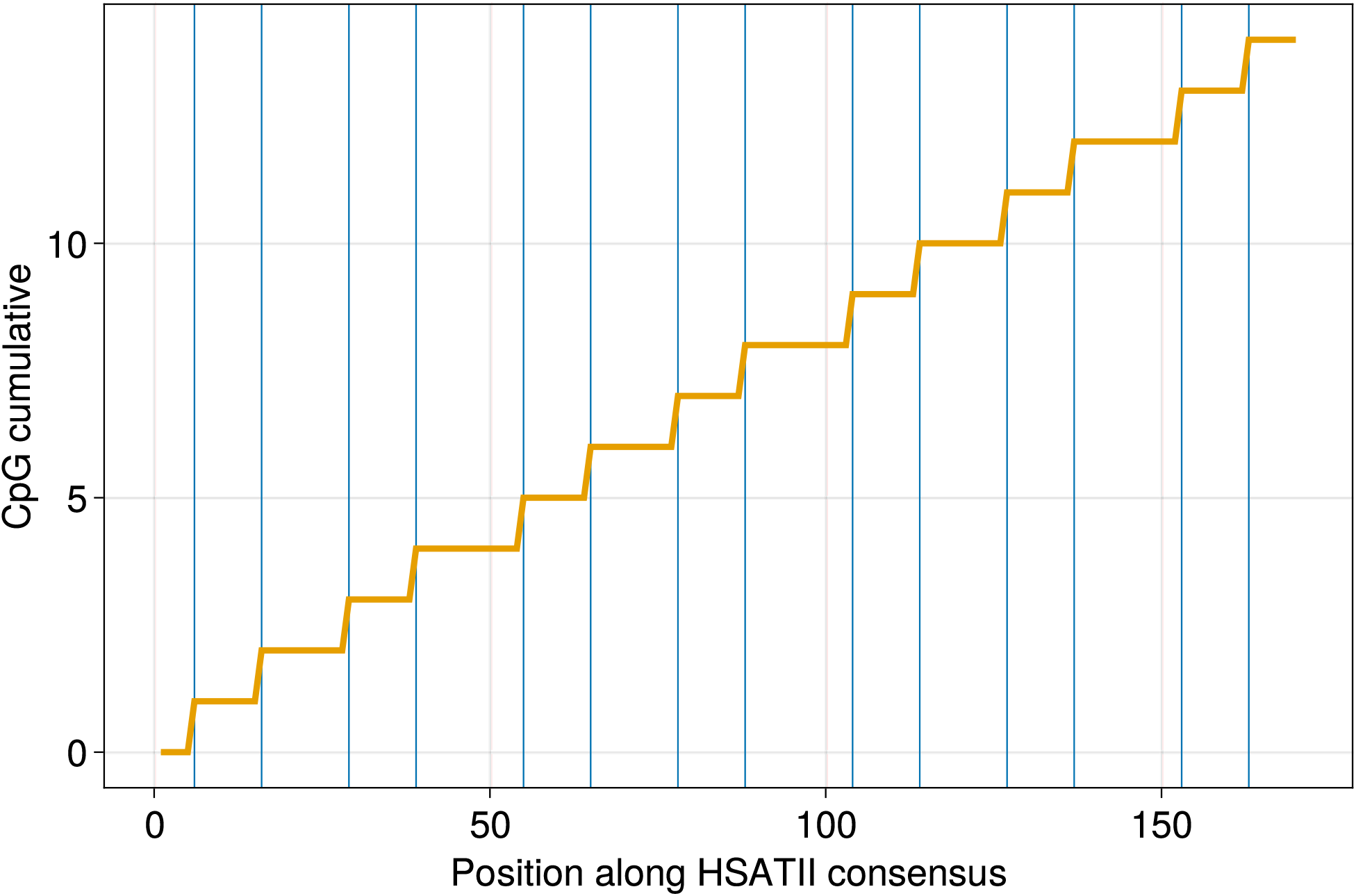
CpG occurrence (vertical blue lines) in the consensus HSATII sequence as annotated in DFAM. The orange line denotes the cumulative number of CpGs.

